## Supplementary Information for "Dual-Resolving of Lipid Positional and Geometric Isomers of C=C Bonds *via* Bifunctional Photocycloaddition-Photoisomerization Reaction System and Mass Spectrometry"

### Materials

Fatty acid standards including FA 14:0, FA 16:1 (9*Z*), FA 16:1 (9*E*), FA 16:0, FA 17:1 (10*Z*), FA 17:1 (10*E*), FA 17:0, FA 18:0, FA 18:1 (9*Z*), FA 18:1 (9*E*), FA 18:1 (11*Z*), FA 18:1 (6*Z*), 18:2 (9*Z*, 11*E*), FA 18:2 (9*Z*, 12*Z*), FA 19:1 (10*Z*), FA 19:1 (10*E*), FA 19:0, FA 20:1 (11*Z*), FA 20:1 (11*E*) and FA 20:4 (5*Z*, 8*Z*, 11*Z*, 14*Z*), were purchased from Sigma-Aldrich. Glycerol phospholipids and glycerides were purchased from Avanti Polar Lipids (Alabaster, AL, USA), including PC 18:1 (9*Z*)/18:1 (9*Z*), PC 18:1 (9*E*)/18:1 (9*E*), PC 18:0/18:2 (9*Z*, 12*Z*), PC 16:0/18:1 (9*Z*), PC 18:1 (9*Z*)/16:0, DG 16:0/18:1 (9*Z*), TG 18:1 (9*Z*)/18:1 (9*Z*)/18:1 (9*Z*), TG 18:2 (9*Z*, 12*Z*)/18:2 (9*Z*, 12*Z*)/18:2 (9*Z*, 12*Z*). The lipid nomenclature introduced by Liebisch et al.[1] is adapted in this study. The notations for double bond positions counted from the carboxyl terminal, where the carbon atom was noted “1”. Consequently, a fatty acid with 18 carbon atoms and one *cis*-C=C between 9th and 10th carbon was abbreviated as FA 18:1 (9*Z*). Methyl benzoylformate and Ir[dFppy]$_2$(dtbbpy)PF$_6$ was purchased from Sigma-Aldrich. Other solvents were purchased from Innochem (Beijing, China) and meet or exceed the analytical grade standard.

### Methods

**Visible-light-activated photocatalytic reaction.** A 40 W Blue LED Lamp (Kessil A160WE Tuna Blue) was used to trigger the photocatalytic reaction. Typically, 100 $\mu$L of reaction solution containing methy benzoylformate (MBF, 100 mM), olefins [lipid standards (5-10 mM) or lipid extract from biological samples] and photocatalyst Ir[dFppy]$_2$(dtbbpy)PF$_6$ (5 mol% of MBF) was prepared in methanol and acetonitrile (1:1, v/v) in a micro volume autosampler vial, and the dissolved oxygen was removed by bubbling nitrogen through a needle of a 2 mL syringe for 30 seconds. Then, the LED light was placed on the side of the glass vial to irradiate the reaction solution for 10 min. The distance between the lamp and the vial was about 8 cm. Two small fans were also used to maintain the reaction solution at room temperature. The solution was directly subjected to mass spectrometric analysis after the reaction except for biological sampels, which was centrifuged at 15,000 rpm for 10 min before the MS ananlysis.

**Preparation of the bacterial samples.** Four bacterial samples were used in this study, including one model pathogen in *Arabidopsis* resistance study (*Pseudomonas syringae*, *P. syringae*) and three endophytic bacteria isolated from *Arabidopsis* leaves (*Weizmannia ginsengihum*,*W. ginsengihum*; *Pseudomonas citronellolis*,*P. citronellolis* and *Moraxella osloensis*,*M. osloensis*). To isolate the endophytic bacteria, *Arabidopsis thaliana* Col-0 plants were grown in unsterilized soil at 22 $^{o}$C, with relative humidity of 60% and photoperiod at a 12:12 h light: dark cycle. 5 weeks later, mature leaves were detached and surface sterilized in 75% alcohol for 90 sec, followed by washing with sterile water for three times. Leaves were then dried on sterile filter papers. Three leaves were put into a 1.5 mL Eppendorf tube filled with 500 $\mu$L 10 mM $MgCl_2$ and one steel ball. Leaves were ground with a grinder. After brief centrifugation, supernatant was plated on R2A plates and bacteria were grown at room temperature for 3 days. About 100 colonies were picked and cultured by R2A liquid medium on a shaker at 220 rpm and 28 $^{o}$C overnight. 16S rRNA fragments were amplified using primers 27F/1492R and then sequenced. Sequence blast identified different endophytic bacteria and three of them were randomly selected for this study.

For the extraction of lipids from these samples, 400 $\mu$L Methyl tert-Butyl Ether (MTBE) was added into the 1.5 mL Eppendorf (EP) tube that contains bacterial cells collected by centrifugation, and subjected to vortex for 30 s and ultrasonic mixing for 10 min in ice water. Then, 80 μL MeOH and 200 μL $H_2O$ were added successively with the same steps as above. After the vortex and sonicate, the mixture was centrifuged for 15 min at 3,000 rpm to separate phases. The upper MTBE phase was taken to a new EP tube and dried with nitrogen flow. The dried lipid extracts were redissolved in methanol and acetonitrile (1:1, v/v) and subjected to photocatalytic reaction.

**Animals.** All experiments and procedures were approved by the committee of experimental animals of Tongji Medical College. Male C57BL/6 mice (wild-type) (8–10weeks old, 23–25 g) were purchased from Wuhan University Laboratory Animal Center. All experimental protocols and animal handling procedures were performed in accordance with the National Institute of Health Guide for the Care and Use of Laboratory Animals (NIH Publications No. 80-23) revised 1996 and the experimental protocols were approved by the committee of experimental animals of Tongji Medical College.

**Middle cerebral artery occlusion (MCAO) model of focal ischemia.** The animals were maintained at temperature of 23±1 °C with 50±10% relative humidity in the Experimental Animal Center. Mice were allowed free access to food and water in a 12-h light/dark cycle. Focal cerebral ischemia was induced by transient occlusion of the right middle cerebral artery (tMCAO) with a 6-0 silicone-coated nylon monofilament, as previously described.[2] Anesthesia was induced with ketamine (100 mg/kg, i.p.) and xylazine (8 mg/kg, i.p.). Focal cerebral ischemia was induced by transient occlusion of the right middle cerebral artery (tMCAO) with a 6-0 silicone-coated nylon monofilament. Briefly, under the operating microscope, the right common carotid artery (CCA), the right external carotid artery (ECA), and the right internal carotid artery (ICA) were isolated and a 6-0 suture was tied at the origin of the ECA and at the distal end of the ECA. The right CCA and ICA were temporarily until resistance was felt and the filament was inserted about 9 to 10 mm from the carotid bifurcation, effectively blocking the middle cerebral artery (MCA). The diameter of the tip of coated suture was considered acceptable between 180 and 220 micrometers. The suture remained inserted for 60 minutes, after which it was removed for reperfusion and the ECA was permanently tied. Body temperature was maintained at 37 ± 0.5°C during and after the surgery with a heating pad. During these experiments, blood pressure (BP) was monitored through a femoral artery and blood gas was analyzed at the end (i.e. 3 days after tMCAO procedure). Subcutaneous normal saline (0.9%) was administered daily, adjusting the volume according to the animal's weight loss.

**Preparation of mouse brain tissues.** For the measurement of lipid changes, C57BL/6 mice were subject to 60 minutes ischemia followed by 1- and 3-days reperfusion. The brains of the mice were collected and cut into two halves from the middle line namely left (normal part) and right half (ischemic part). Then the cut parts were weighed and added into three-times of methanol [weight (mg) to volume (mL)] for homogenization at 40 Hz for 2 min. The mixture was then diluted 30 times with methanol solution. The mixed methanol solution was divided into 300 μL fractions for further lipid extraction. Briefly, 300 μL of homogenate in methanol were added into 1 mL MTBE and vortexed for 10 s. Subsequently, the mixture was vibrated for 10 min. Then 300 μL of water was added and vortexed for 10 s to promote liquid-liquid stratification. After equilibration for 10 min, the mixture was centrifuged at 12,000 rpm for 10 min at 4 °C. A total of 900 μL of organic phase was transferred to a new tube for dry with nitrogen flow. The dried lipid extracts were reconstituted in 100 $\mu$L (equivalent of 10 mg original tissue samples) solution of methanol and acetonitrile (1:1, v/v), and subjected to photocatalytic reaction and LC-MS analysis.

**Liquid chromatographic and mass spectrometric analysis.** The data for reaction method optimization and relative quantitation of C=C location isomer were accomplished with LTQ-Orbitrap Elite mass spectrometer (Thermo Scientific, Germany). The main mass parameters were set up as below: sheath gas ($N_2$): 40 arbitrary units; auxiliary gas ($N_2$): 5 units; capillary temperature: 350 $^{o}$C; spray voltage: 3.8 kV; collision energy: 40 V; the resolution of full mass scan, 30,000. The parameters of MS/MS scan were set as below: top 5 ions with high intensity were selected for fragmantation analysis, and dynamic exclusion function was turned on for 10 s. The MS/MS scan was induced by collision dissociation (CID), with a resolution of 15,000.

The analysis for C=C location and bacteria cells samples was performed on a hybrid trapped ion mobility-quadrupole time-of-flight mass spectrometer (timsTOF Pro, Bruker Daltonics, Bremen, Germany). This QToF instruments were operated to collect full scan MS data and MS/MS fragmentation spectra in the same analytical run with the parallel accumulation serial fragmentation (PASEF) method. Precursors for data-dependent acquisition were isolated within ± 1 Th and fragmented with an ion mobility-dependent collision energy, which was linearly increased from 25 to 45 eV in positive mode, and from 35 to 55 eV in negative mode. The ion mobility was scanned from 0.6 to 1.95 Vs/cm$^2$. Low-abundance precursor ions with an intensity above a threshold of 100 counts but below a target value of 4000 counts were repeatedly scheduled and otherwise dynamically excluded for 0.2 min. TIMS ion charge control was set to 5e6. The TIMS dimension was calibrated linearly using four selected ions from the Agilent ESI LC/MS tuning mix. The ion source settings were: capillary voltage = 4.5 kV; end plate offset = 500 V; drying gas flow = 12.0 L/min; nebulizer gas = 5.0 bar; drying temperature = 250 °C. The data acquisition rate was set to 8 Hz over the mass range of *m/z* 30–1000. The sodium adducts were used for fragmentation in CID to identify the C=C positional isomers of fatty acids, as both the deprotonated and protonated ions could not generate the anticipated diagnostic ions. For DG and TG, ammonium adducts were used for fragmentation to determine the isomers. Other glycerol phospholipids were fragmented in protonated forms to generate the diagnostic ions.

A Waters ACQUITY UPLC BEH C18 Column (2.1 mm × 100 mm, 1.7 mm) was performed by the UltiMate 3000 UPLC system (DIONEX, Thermo Scientific, Germany) and maintained at 40 $^{o}$C for separation of lipid species. The flow rate was set as 0.3 $\mu$L/min. Gradient elution consisted of mobile phase A [(ACN/$H_2O$, 60:40, v/v) mixed with 10 mM $NH_4OAc$ and 0.1% formic acid] and mobile phase B [(IPA/ACN, 90:10, v/v) mixed with 10 mM $NH_4OAc$ and 0.1% formic acid]. The optimal chromatographic gradient program for biological samples and GPLs standards was as follow: 30% B at 0~4 min, 30~52% B at 4~6 min, 52~63% B at 6~8 min, 63~68% B at 8~9 min, 68~74% B at 9~27 min, 74~80% B at 27~29 min, 80~99% B at 29~31 min, 99% B at 31~35 min; For FA, the The optimal chromatographic gradient program were shorten as below: FA 30% B at 0~4 min, 30~52% B at 4~6 min, 52~63% B at 6~8 min, 63~68% B at 8~10 min.

**Gas chromatography mass spectrometric analysis.** Gas chromatography mass spectrometric (GC-MS) analysis was carried out by a gas chromatograph (Agilent GC-2030) coupled to a mass spectrometer (Agilent GCMS-QP2020 NX), using a Rtx-5MS fused silica column (30 m × 0.25 mm, 0.25 μm film thickness). The experiments were performed according to the following conditions: injector temperature, 280 °C; injection in spitless mode; gas flow, 0.6 mL/min; and injection volume, 8 μL. The oven temperature was programmed as follows: initial temperature 140 °C, hold for 1 min; ramp at 10 °C/min up to 180 °C, hold for 20 min, then ramp at 0.5 °C/min up to 200 °C, hold for 20 min, finally ramp at 5 °C/min up to 230 °C, hold for 2 min. The total analysis time was 89 min. The single quadrupole mass spectrometer was operated in the full scan mode, with the instrumental temperatures set at 250, 250, and 180 °C for transfer line, source, and quadrupole, respectively. The electron energy was set at 70 eV, data acquisition was carried out in an *m/z* range from 45 to 550 and with a solvent delay for 4.5 min.

The free fatty acids were methylated with 3 mL 0.5 mol/L $H_2SO_4$-MeOH at 90 °C for 60 min. After the reaction, 3 mL 0.4 mol/L NaOH-MeOH was added for neutralization of the solutions. Then, 5 mL saturated normal saline and 2 mL hexane were added in sequentially for promoting liquid-liquid stratification and extracting the methylated fatty acids. The hexane phase was transferred into a new centrifuge tube for drying with anhydrous sodium sulfate. The obtained solution was concentrated with nitrogen blowing. Finally, the residual was redissolved with 200 uL hexane and centrifuged at 10000 rpm for 10 min for the analysis of GC-MS.

**Data analysis.** Thermo Xcalibur mass spectrometry data system was used for extract target ion and calculate its peak area, namely quantitation analysis. MS-DIAL version 4.12 was used for the process of LC-MS data for qualitative analysis. The raw data must be transformed into Abf. files. More details were presented at http://prime.psc.riken.jp/compms/msdial/main.html. The acquired GC-MS data were processed with GCMS solution Version 5.2 (Agilent, USA). The mass spectra matching was processed with a standard Library NIST11. In focal ischemia and reperfusion experiments, six of biological replicates were conducted for 1-day and 3-day reperfusion, respectively. Two-tailed Student's t-test was used to compare the differences of the ratios of lipid positional isomers between 1-day and 3-day reperfusion.

**Computational methods.** All density functional theory (DFT) calculations were carried out using the Gaussian 16 software package.[3] The geometries were optimized using the M06-2X functional[2] with a basis set of 6-31G(d) for all atoms. Vibrational frequency calculations were performed for all the stationary points to confirm if each optimized structure is a local minimum or a transition state structure. Truhlar's quasi-harmonic corrections[3] were applied for entropy calculations with 100 $cm^{-1}$ as the frequency cutoff using the Goodvibes program.[4] Intrinsic reaction coordinate (IRC) calculations were performed to ensure that the saddle points located were transition states connecting the reactants and the products. Solvation energy corrections were calculated in acetonitrile solvent with the SMD continuum solvation model[5] based on the gas phase optimized geometries. The M06-2X functional with a basis set of 6-311+G(d,p) for all atoms was used for single-point energy calculations.

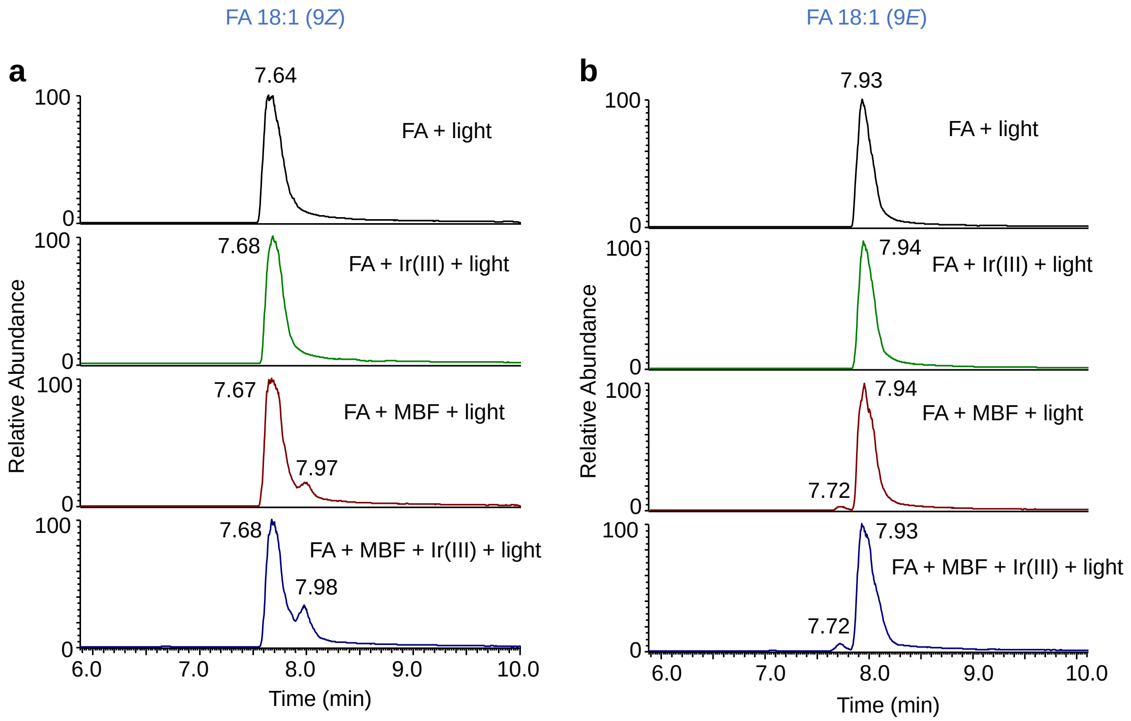


**Figure S1.** EICs of FA 18:1 at different reaction conditions. (a)-(b) EICs of (a) FA 18:1 (9*Z*) and (b) FA 18:1 (9*E*) at reaction conditions of only LED (450 nm) light irradiation, light irradiation in the presence of Ir[dFppy]$_2$(dtbbpy)PF$_6$ (Ir(III)) photocatalyst, light irradiation in the presence of methyl benzoylformate (MBF) and light irradiation in the presence of both MBF and Ir(III) photocatalyst. Reaction time: 10 min.

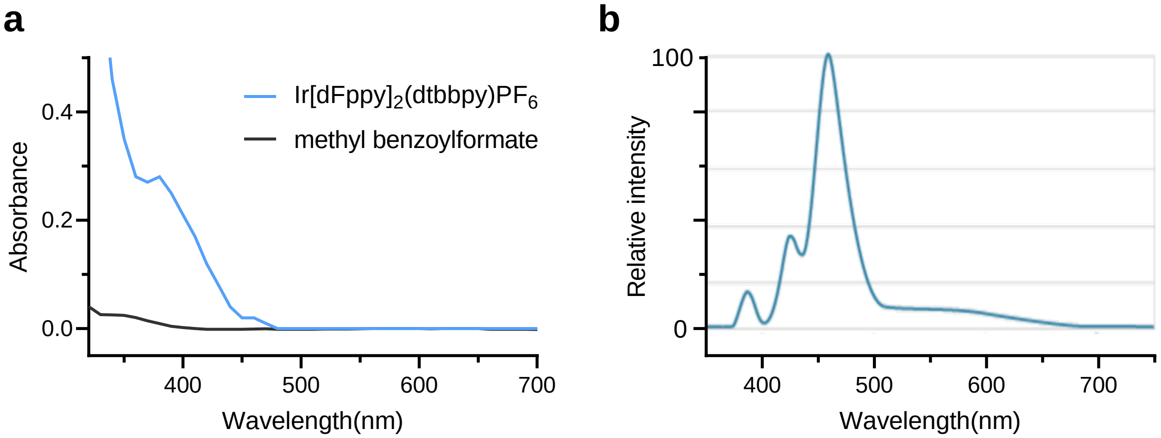


**Figure S2**. Spectroscopic characterization of the reactant, photocatalyst and LED light. (a) Ultraviolet-visible absorption spectra of the solutions of photocatalyst Ir[dFppy]$_2$(dtbbpy)PF$_6$ (0.1 mM) and methyl benzoylformate (1 mM) in MeOH. (b) Full spectrum of the Kessil A160WE-TB blue LED light adopted from vendor's website.

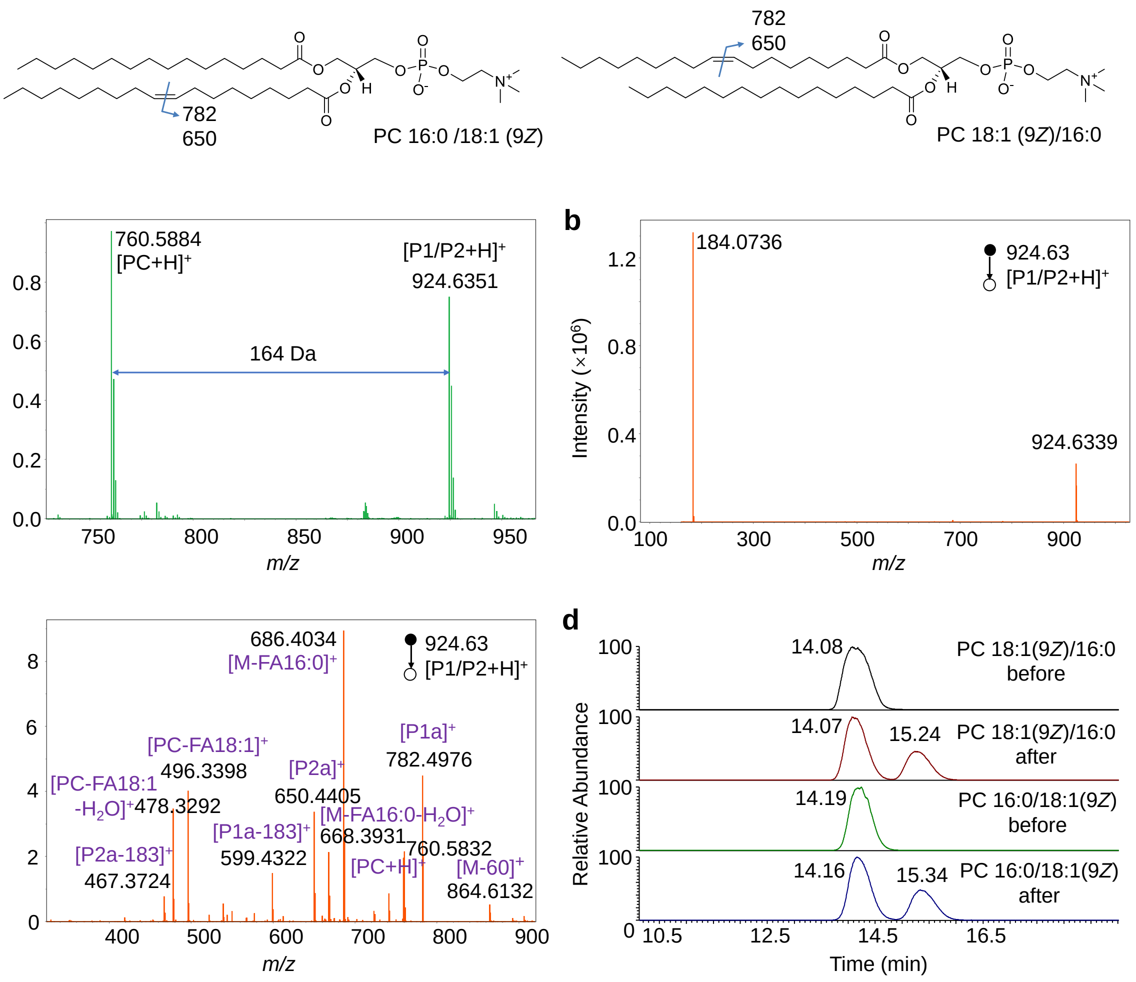


**Figure S3**. Demonstration of the bifunctional reaction system for the identification of C=C bonds location and *E*-*Z* configuration in PC 16:0/18:1 (9*Z*) and 18:1 (9*Z*)/16:0. (a) Mass spectrum of the photocatalytic reaction solution of PC 16:0/18:1 (9*Z*) in positive ion mode. (b)-(c) MS/MS spectrum of the protonated photocycloaddition product P1/P2 in positive ion mode in the mass range of (b) 100-1000 Da and (c) 200-900 Da. (d) Extracted ion chromatograms (EIC) of PC 18:1 (9*Z*)/16:0 and 16:0/18:1 (9*Z*) before and after the photocatalytic reaction in positive ion mode.

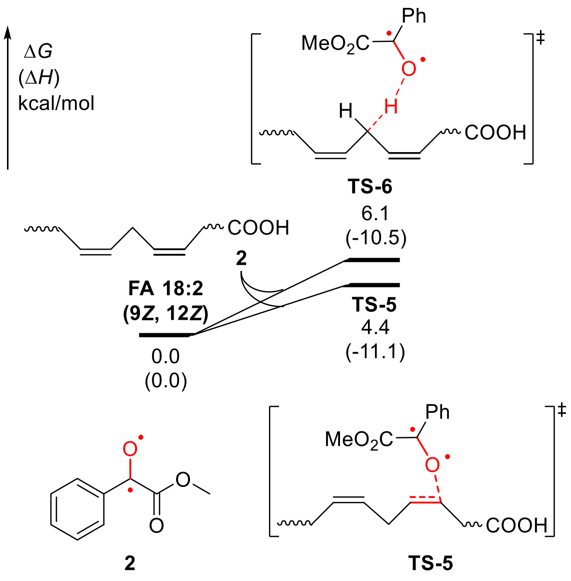


**Figure S4.** Computational study of the comparison between hydrogen atom abstraction and radical addition to alkene in the case of FA18:2 (9*Z*, 12*Z*).

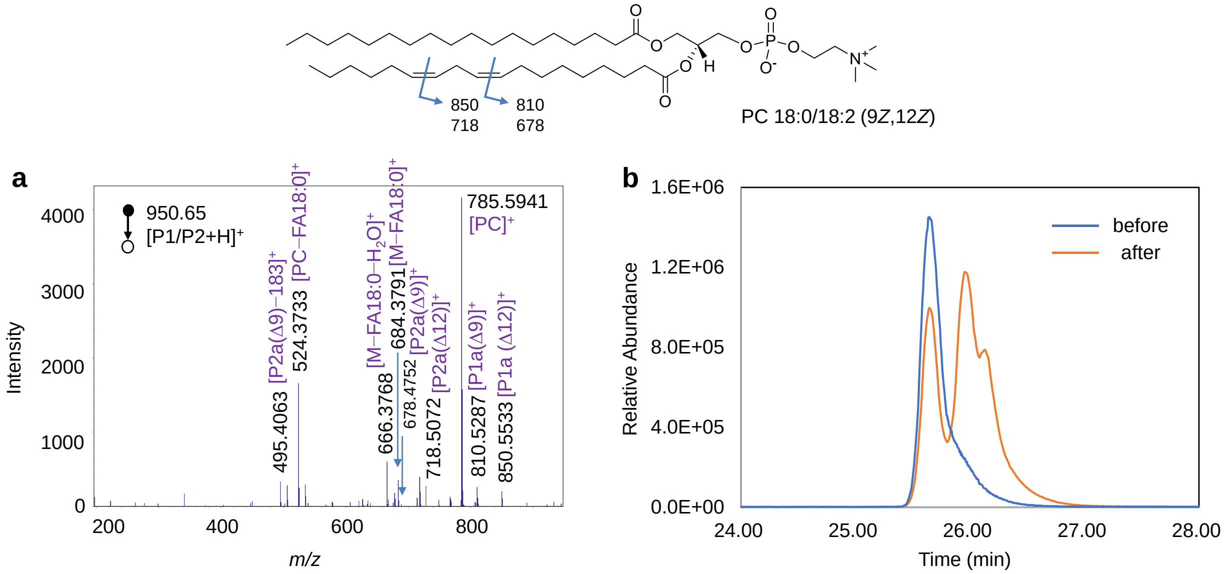


**Figure S5.** Demonstration of the bifunctional reaction system for the identification of C=C bonds location and *E-Z* configuration in PC 18:0/18:2 (9*Z*, 12*Z*). (a) MS/MS spectrum of the protonated photocycloaddition product (P1/P2) of PC 18:0/18:2 (9*Z*, 12*Z*). (b) EICs of PC 18:0/18:2 (9*Z*, 12*Z*) before and after the photocatalytic reaction at *m/z* 786.61.

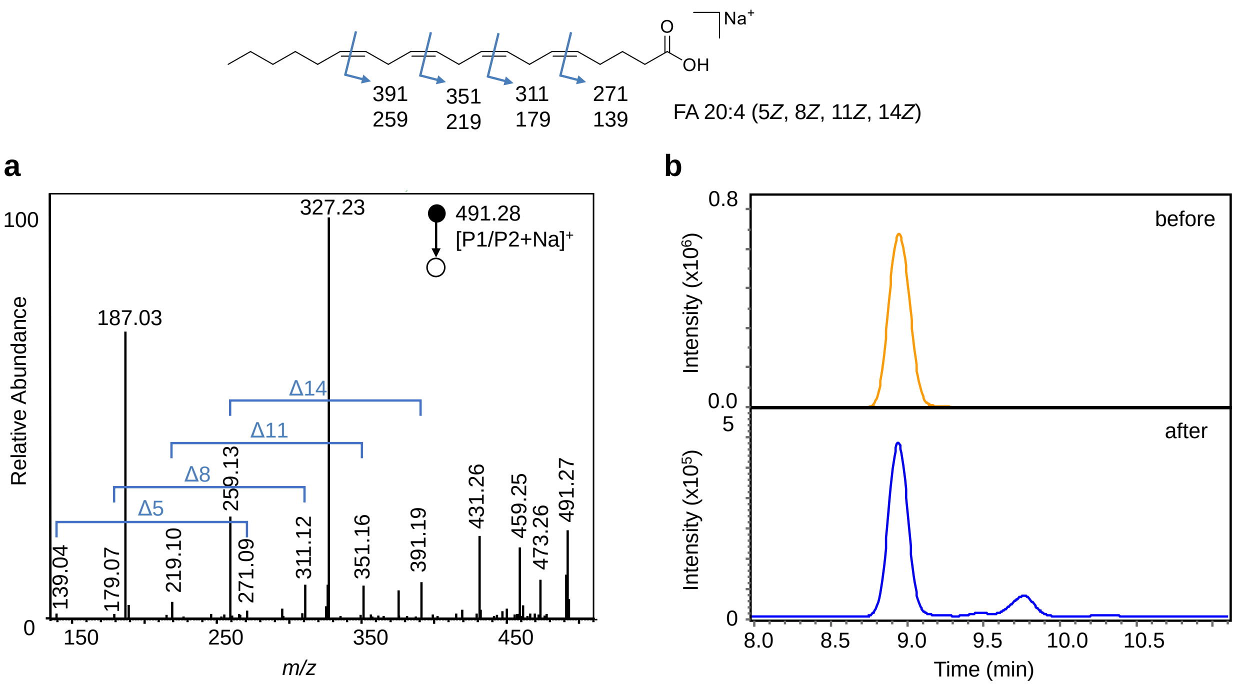


**Figure S6.** Demonstration of the bifunctional reaction system for the identification of C=C bonds location and *E*-*Z* configuration in FA 20:4 (5*Z*, 8*Z*, 11*Z*, 14*Z*). (a) MS/MS spectrum of the sodiated photocycloaddition product (P1/P2) at *m/z* 491.28. (b) EICs of FA 20:4 (5*Z*, 8*Z*, 11*Z*, 14*Z*) before and after the photocatalytic reaction at *m/z* 467.28 in negative ion mode.

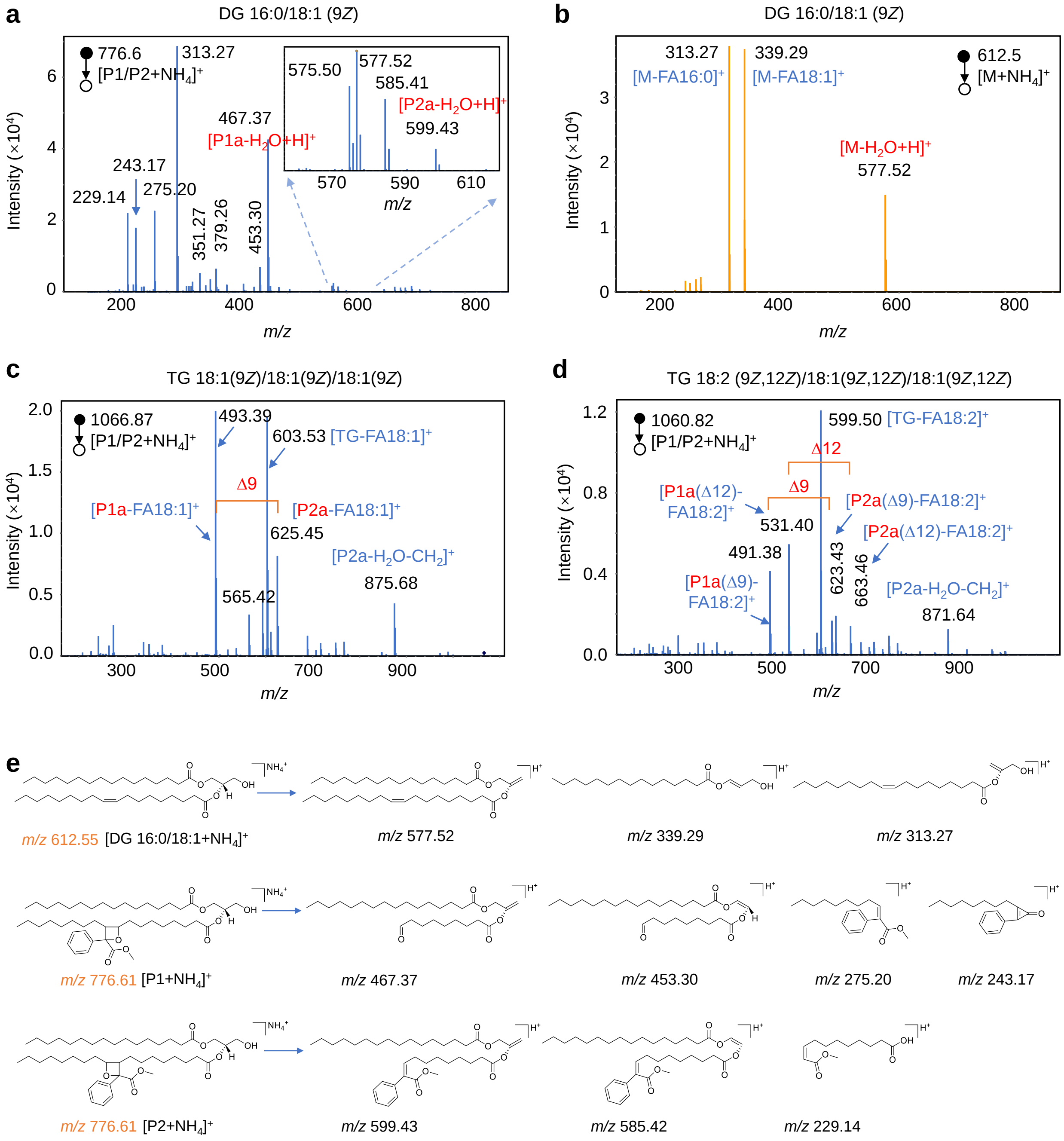


**Figure S7.** Identification of C=C bonds in glycerides. (a)-(d) MS/MS spectrum of the photocycloaddition products of (a) DG 16:0/18:1 (9*Z*), (c) TG 18:1 (9*Z*)/18:1 (9*Z*)/18:1 (9*Z*), (d) TG 18:2 (9*Z*,12*Z*)/18:2 (9*Z*,12*Z*)/18:2 (9*Z*,12*Z*), and (b) MS/MS spectrum of DG 16:0/18:1 (9*Z*) before the reaction. (e) Possible fragmentation pathways for DG 16:0/18:1 (9*Z*) and its cycloaddition products.

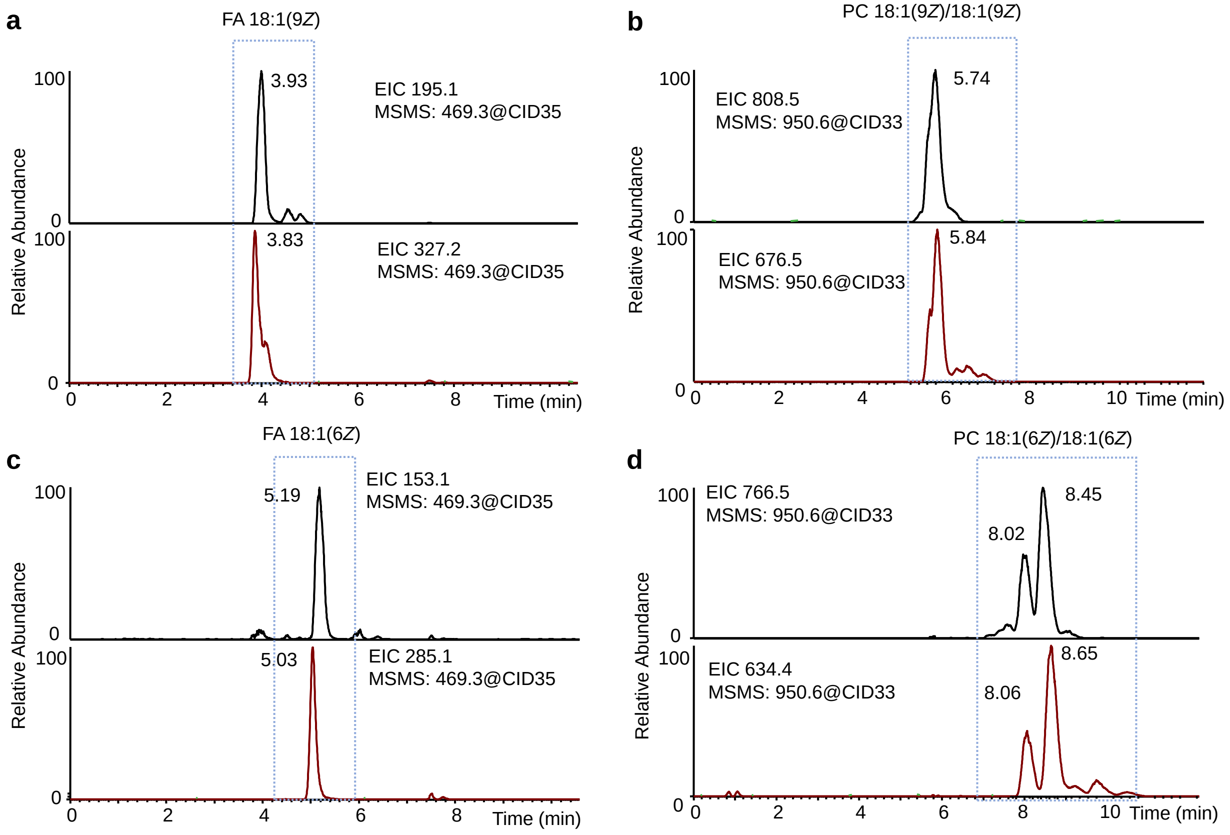


**Figure S8.** EICs of diagnostic ions in tandem MS spectra of photocycloaddition products of (a) FA 18:1 (9*Z*), (b) PC 18:1 (9*Z*)/18:1 (9*Z*), (c) FA 18:1 (6*Z*), and (d) PC 18:1 (6*Z*)/18:1 (6*Z*) in the positive ion mode for quantitative analysis.

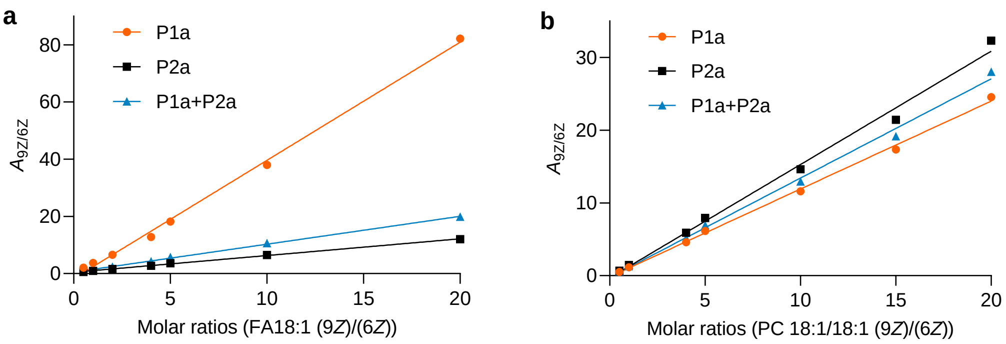


P1a: $y = 0.5855x + 0.4519$, $R^2 = 0.9986$
P2a: $y = 4.1312x - 1.7198$, $R^2 = 0.9973$
P1a+P2a: $y = 0.9755x + 0.5144$, $R^2 = 0.9987$

P1a: $y = 1.5582x - 0.3106$, $R^2 = 0.9931$
P2a: $y = 1.2078x - 0.1689$, $R^2 = 0.9981$
P1a+P2a: $y = 1.3646x - 0.2386$, $R^2 = 0.9959$

**Figure S9.** Quantitative analysis of the C=C positional isomers. (a) Linear relationship established between the peak area ratio ($A_{9Z}/A_{6Z}$) of the diagnostic ions and molar ratio ($C_{9Z}/C_{6Z}$) of the two FA 18:1 C=C location isomers. (b) Linear relationship established between the peak area ratio ($A_{9Z}/A_{6Z}$) of the diagnostic ions and molar ratio ($C_{9Z}/C_{6Z}$) of the two PC 18:1/18:1 C=C location isomers.

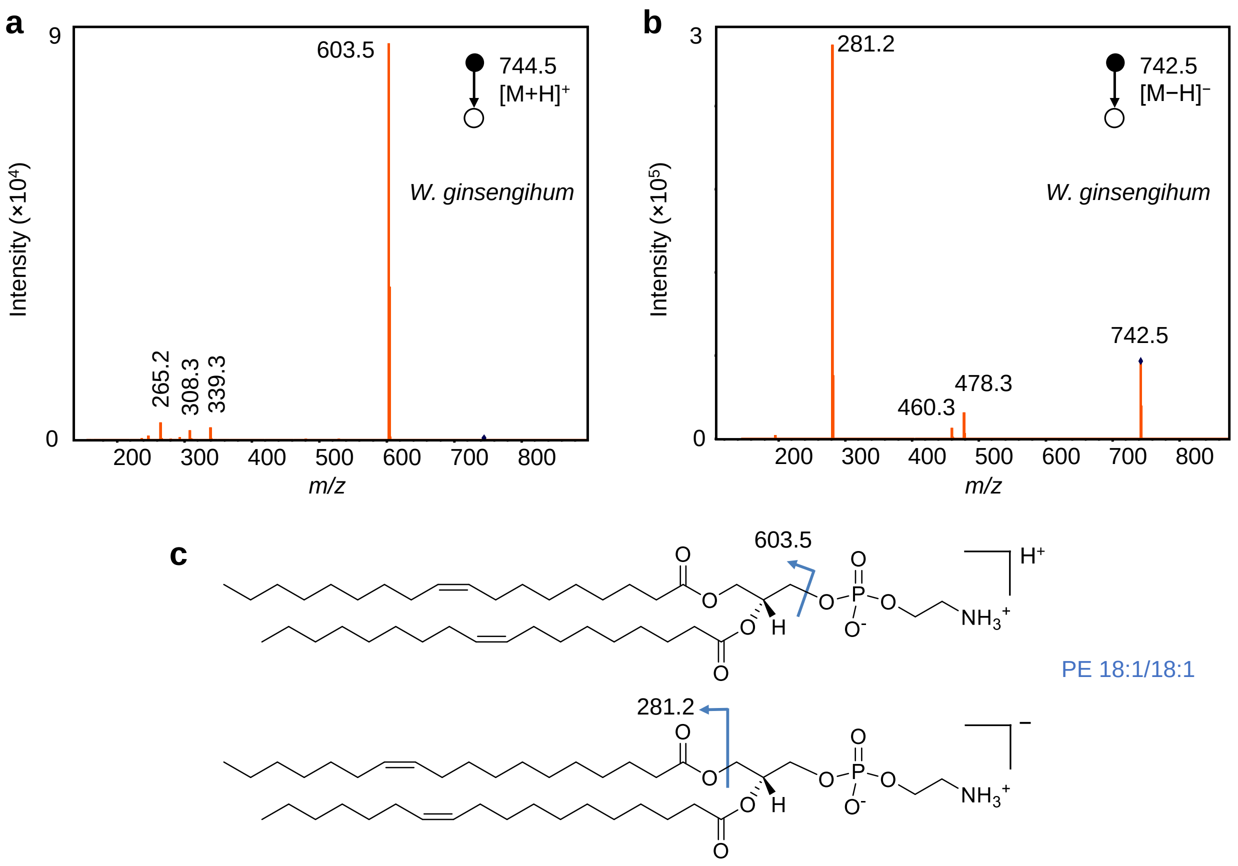


**Figure S10**. Analysis of the structure of PE 36:2 by tandem MS. (a) MS/MS spectrum of $[M+H]^+$ at *m/z* 744.5 in positive ion mode. (b) MS/MS spectrum of $[M-H]^-$ at *m/z* 742.5 in negative ion mode. (c) Structures of PE 18:1 ($\Delta$9)_18:1 ($\Delta$9) ions and possible fragmentation pathways in positive and negative ion modes, respectively.

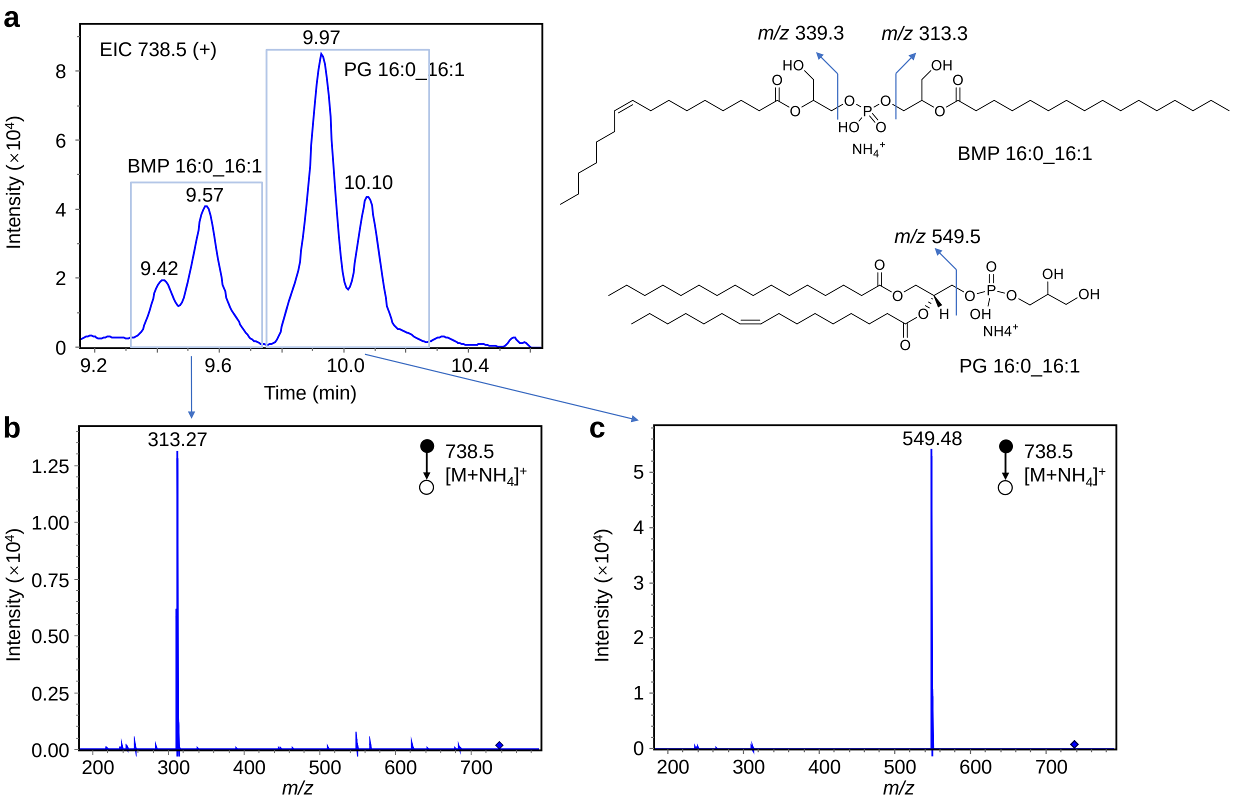


**Figure S11.** Identification of bis(monoacylglycero)phosphate (BMP) in *P. citronellolis* sample. (a) EIC of *m/z* 738.5 that shows the presence of BMP 32:1 and PG 32:1 isomers. (b) MS/MS spectrum of $[M+NH_4]^+$ ion for BMP 32:1. (c) MS/MS spectrum of $[M+NH_4]^+$ ion for PG 32:1. The different characteristic fragment ions at *m/z* 313.27 and 549.48 indicate the structures of BMP and PG, respectively.

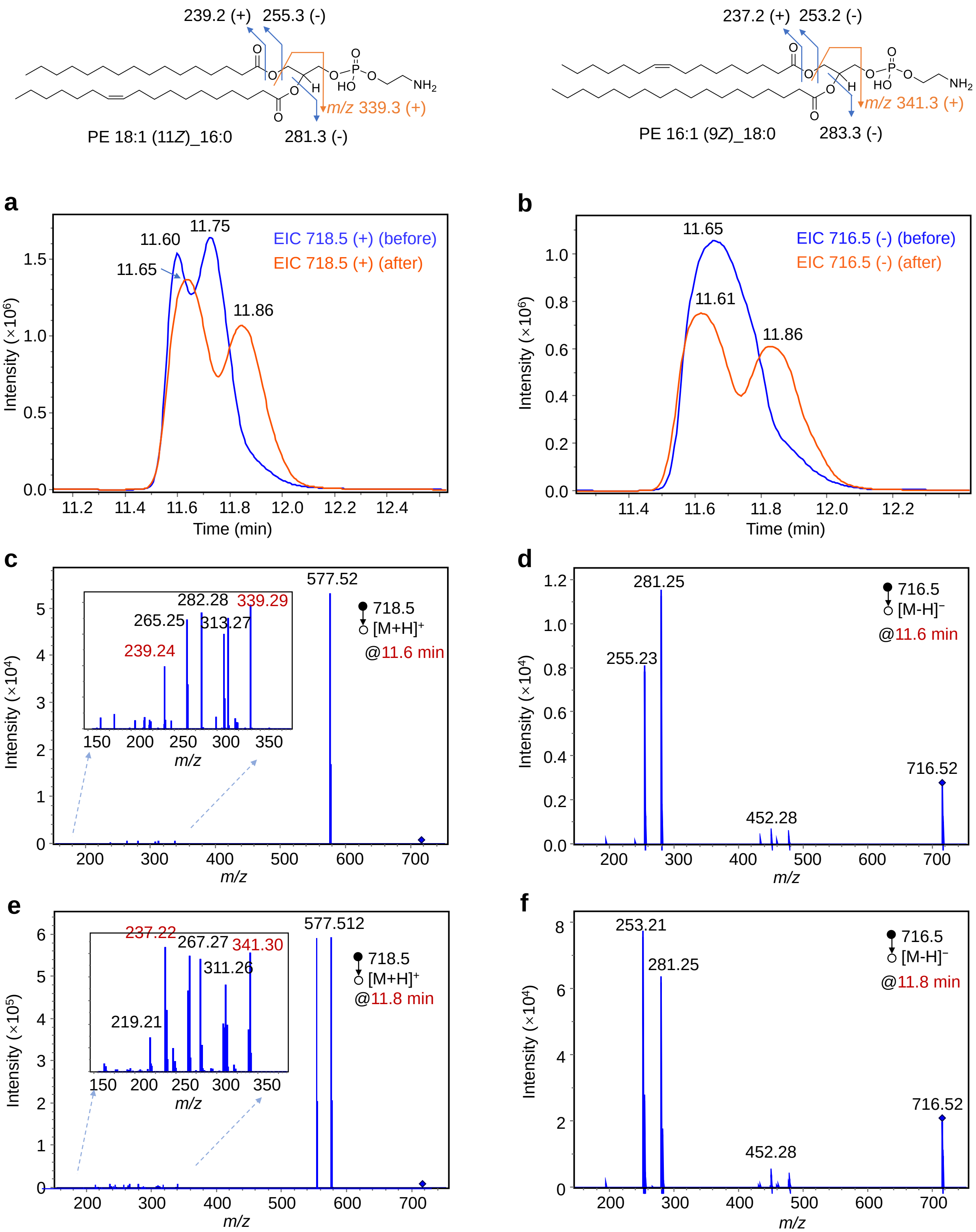
239.2 (+)
255.3 (-)
m/z 339.3 (+)
PE 18:1 (11Z)_16:0
281.3 (-)
237.2 (+)
253.2 (-)
m/z 341.3 (+)
PE 16:1 (9Z)_18:0
283.3 (-)
a
EIC 718.5 (+) (before)
EIC 718.5 (+) (after)
11.60
11.75
11.65
11.86
Intensity (×10⁶)
Time (min)
b
EIC 716.5 (-) (before)
EIC 716.5 (-) (after)
11.65
11.61
11.86
c
577.52
282.28
339.29
265.25
313.27
239.24
718.5
[M+H]+
@11.6 min
Intensity (×10⁴)
m/z
d
281.25
255.23
716.52
452.28
716.5
[M-H]−
@11.6 min
e
577.512
237.22
267.27
341.30
311.26
219.21
718.5
[M+H]+
@11.8 min
Intensity (×10⁵)
f
253.21
281.25
716.52
452.28
716.5
[M-H]−
@11.8 min

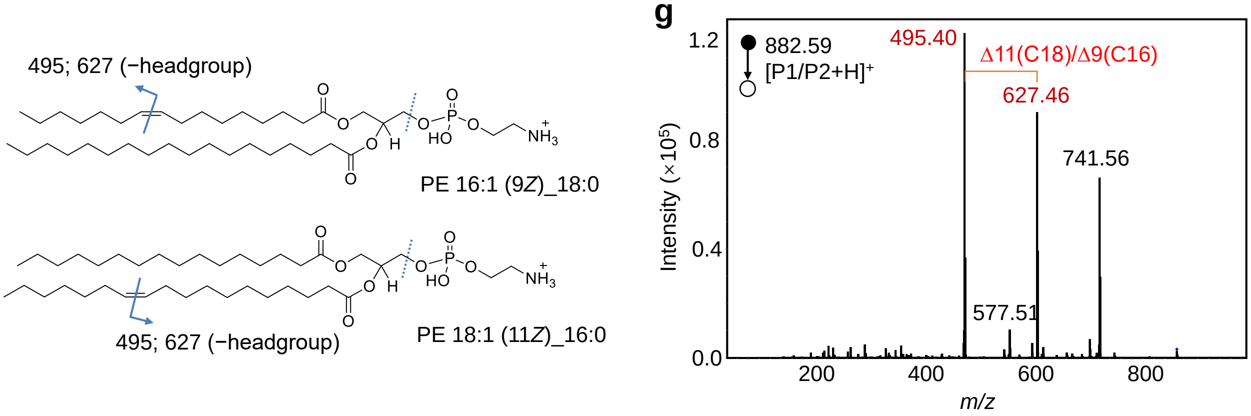


**Figure S12.** Analysis of the structure of PE 34:1 in *P. syringae* bacterial sample. (a) EICs of the protonated ions of PE 34:1 in positive ion mode before and after the photocatalytic reaction. (b) EICs of the deprotonated ions of PE 34:1 in negative ion mode before and after the photocatalytic reaction. The new peak appeared on the right indicated the *cis*-to-*trans* conversion, and confirmed the *Z*-configuration of C=C in these lipids. (c)-(d) MS/MS spectra of (c) $[M+H]^+$ and (d) $[M-H]^-$ for PE 34:1 at retention time of 11.6 min. (e)-(f) MS/MS spectra of (c) $[M+H]^+$ and (d) $[M-H]^-$ for PE 34:1 at retention time of 11.8 min. The neutral loss of 141 Da corresponds to the ethanolamine headgroup indicates the PE subclass of these lipids. The fatty acyl fragments in negative ion modes confirmed the structures of PE 16:1_18:0 and PE 18:1_16:0. (g) Average MS/MS spectrum of the photocycloaddition products. The diagnostic ion pairs at *m/z* 495.40 and 627.46 indicated the isomers of C=C at Δ9 and Δ11, which correspond to the lipid structures of PE 16:1 (9*Z*)_18:0 and PE 18:1 (11*Z*)_16:0. Peak at *m/z* 741.56 corresponds to the fragment of losing ethanolamine headgroup.

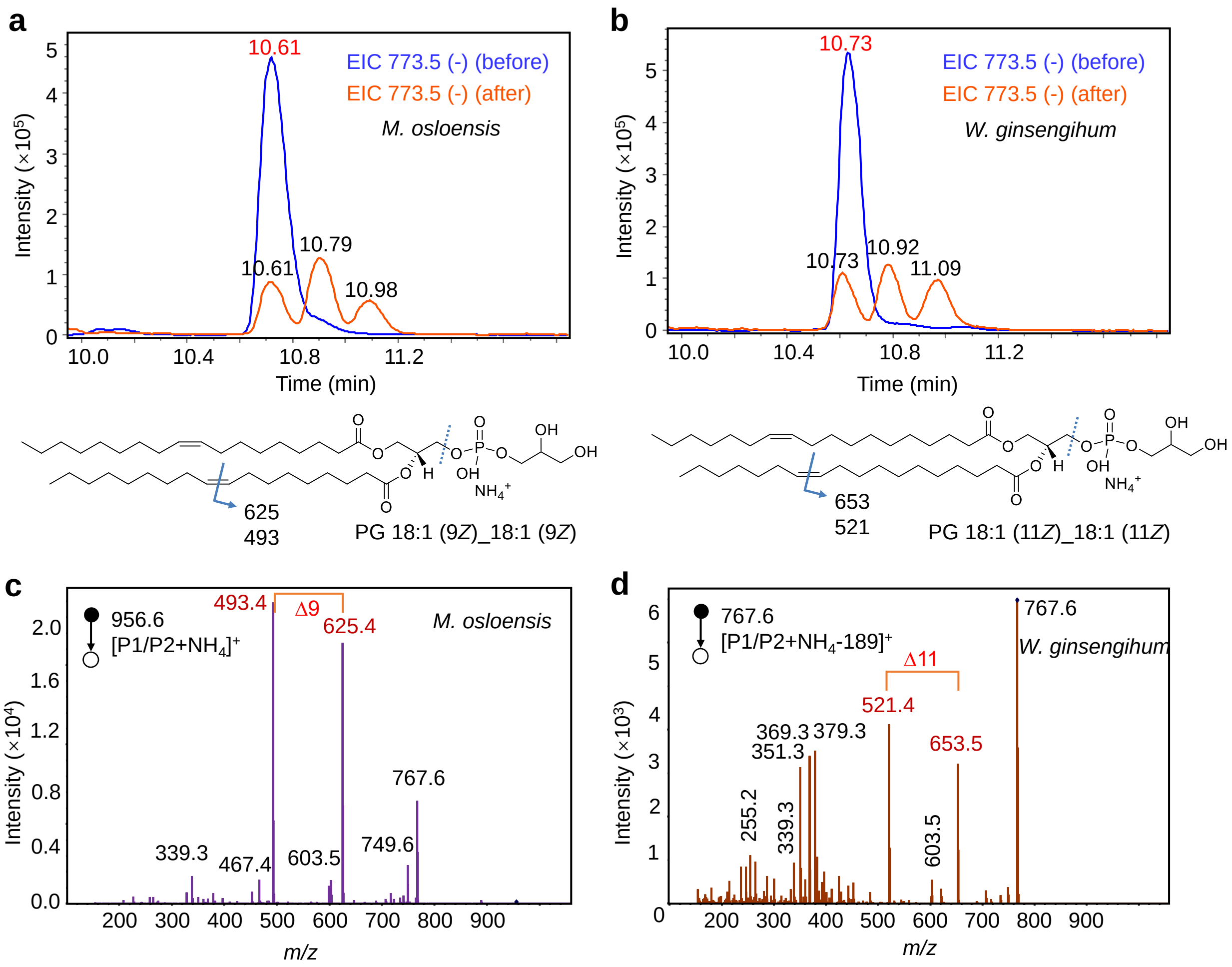


**Figure S13.** Analysis of the structure of PG 36:2 in *M. osloensis* and *W. ginsengihum* bacterial samples. (a)-(b) EICs of the protonated ions of *m/z* 773.5 in negative ion mode before and after the photocatalytic reaction of (a) *M. osloensis* and (b) *W. ginsengihum* samples. The new peak appeared on the right indicated the *cis*-to-*trans* conversion, and confirmed the *Z*-configuration of C=C in these lipids. (c) MS/MS spectrum of the photocycloaddition products of PG 36:2 in *M. osloensis* sample. (d) MS/MS spectrum of ion at *m/z* 767.6, which corresponds to the in-source decay fragment of the photocycloaddition products of PG 36:2 in *W. ginsengihum* sample by losing the head group. The initial formula PG 18:1_18:1 was deduced by the tandem MS spectra in positive and negative ion modes. Based on the diagnostic ion pairs in both samples, only one positional isomer was observed for fatty acyl chains in PG 18:1_18:1. Therefore, the structures were confirmed as PG 18:1 (9*Z*)_18:1 (9*Z*) and PG 18:1 (11*Z*)_18:1 (11*Z*) in *M. osloensis* and *W. ginsengihum*, respectively.

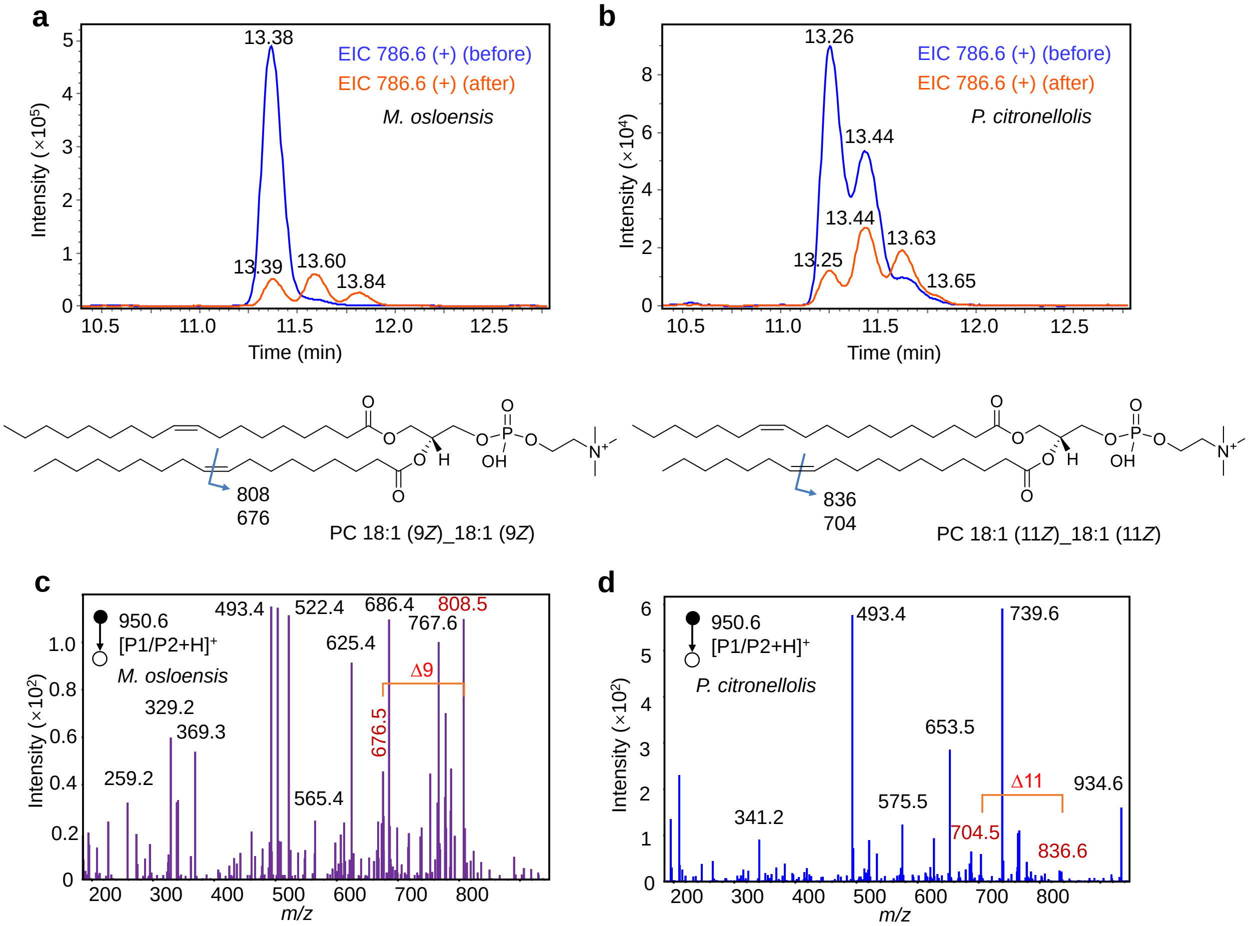


**Figure S14.** Analysis of the structure of PC 36:2 in *M. osloensis* and *P. citronellolis* bacterial samples. (a)-(b) EICs of the protonated ions of *m/z* 786.6 in positive ion mode before and after the photocatalytic reaction of (a) *M. osloensis* and (b) *P. citronellolis* samples. The new peak appeared on the right indicated the *cis*-to-*trans* conversion, and confirmed the *Z*-configuration C=C bond of PC 18:1_18:1 in *M. osloensis*, and both *Z*- and *Z,E*-configuration of C=C bonds of PC 18:1_18:1 in *P. citronellolis* samples, respectively. (c)-(d) MS/MS spectra of the photocycloaddition products of PC 36:2 in (c) *M. osloensis* and (d) *P. citronellolis* samples*.* The initial formula PC 18:1_18:1 was deduced by the tandem MS spectra in positive and negative ion modes. Based on the diagnostic ion pairs in both samples, the structures were confirmed as PC 18:1 (9*Z*)_18:1 (9*Z*) in *M. osloensis*, and both PC 18:1 (9*Z*)_18:1 (9*Z*) and PC 18:1 (9*Z*)_18:1 (9*E*) in *P. citronellolis*.

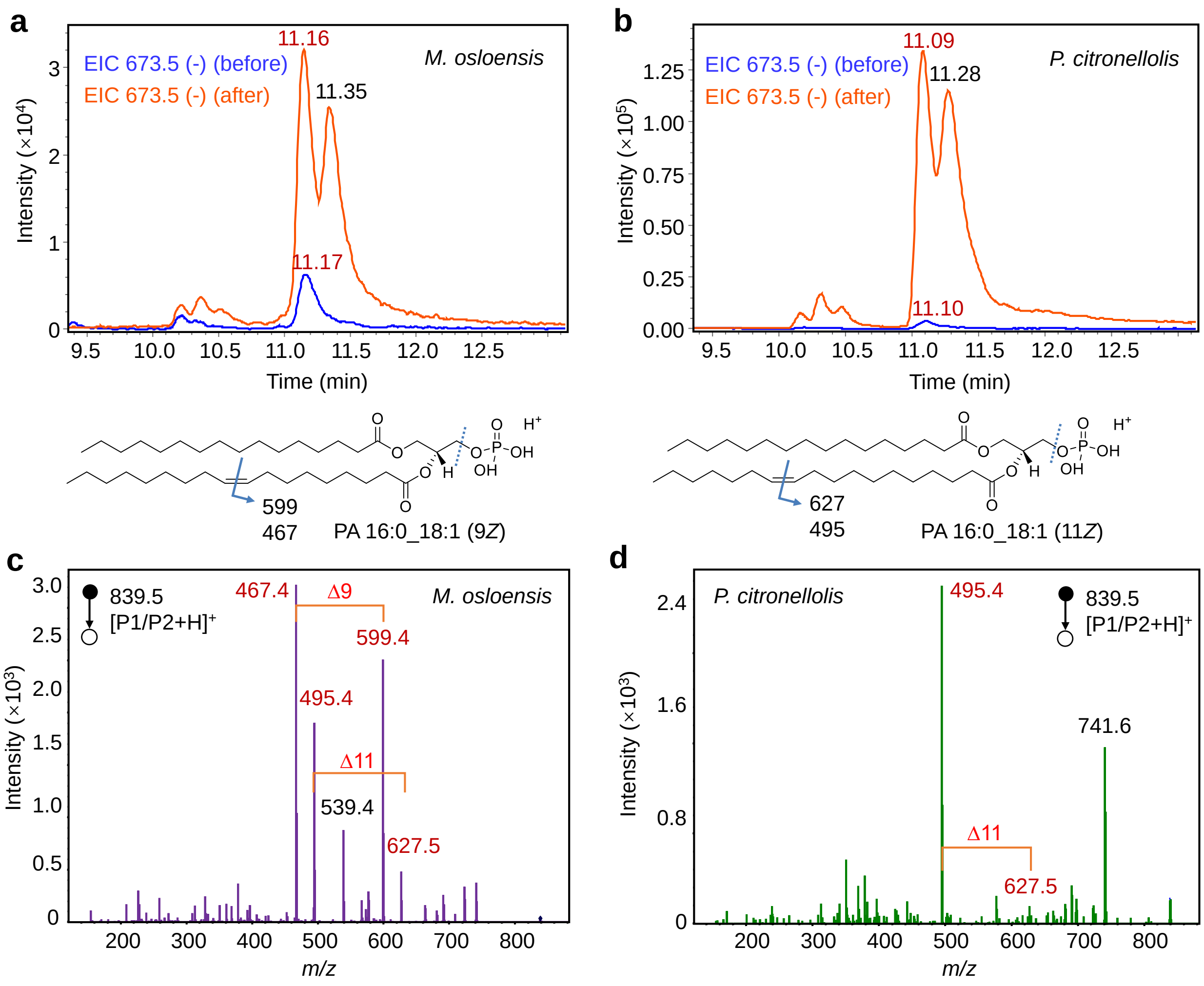


**Figure S15.** Analysis of the structure of PA 34:1 in *M. osloensis* and *P. citronellolis* bacterial samples. (a)-(b) EICs of the protonated ions of *m/z* 673.5 in negative ion mode before and after the photocatalytic reaction of (a) *M. osloensis* and (b) *P. citronellolis* samples. The new peak appeared on the right indicated the *cis*-to-*trans* conversion, and confirmed the *Z*-configuration C=C bonds of PA 16:0_18:1 in both *M. osloensis* and *P. citronellolis* samples. (c)-(d) MS/MS spectra of the photocycloaddition products of PA 34:1 in (c) *M. osloensis* and (d) *P. citronellolis* samples. The initial formula PC 16:0_18:1 was deduced by the tandem MS spectra in positive and negative ion modes. Based on the diagnostic ion pairs in both samples, the structures were confirmed as PA 16:0 _18:1 (9*Z*) and PA 16:0 _18:1 (11*Z*) in *M. osloensis*, and PA 16:0 _18:1 (11*Z*) in *P. citronellolis*.

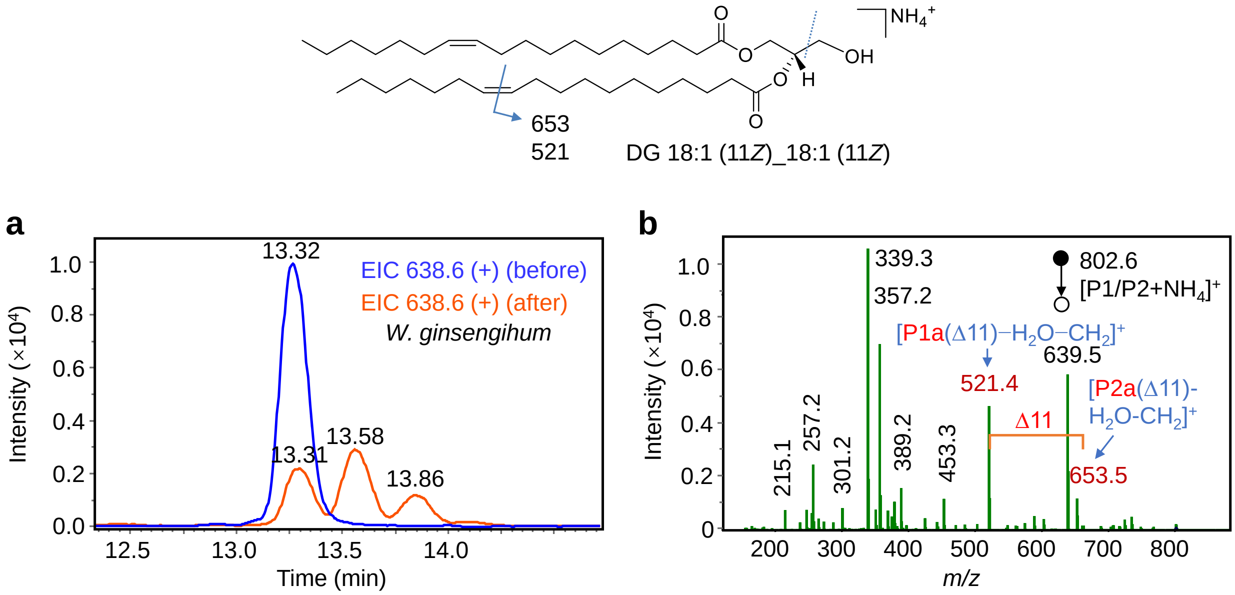


**Figure S16.** Analysis of the structure of DG 36:2 in *W. ginsengihum* bacterial sample. (a) EICs of the ions of *m/z* 638.6 in positive ion mode before and after the photocatalytic reaction. The two new peaks appeared on the right indicated the *cis*-to-*trans* conversion, and confirmed the *Z*-configuration C=C bonds of DG 18:1_18:1. (b) MS/MS spectrum of the photocycloaddition products of DG 36:2 in *W. ginsengihum* sample. The initial formula DG 18:1_18:1 was deduced by the tandem MS spectra in positive and negative ion modes. Based on the diagnostic ion pairs in the MS/MS spectrum of the photocycloaddition products, the structures were confirmed as DG 18:1 (11*Z*)_18:1 (11*Z*).

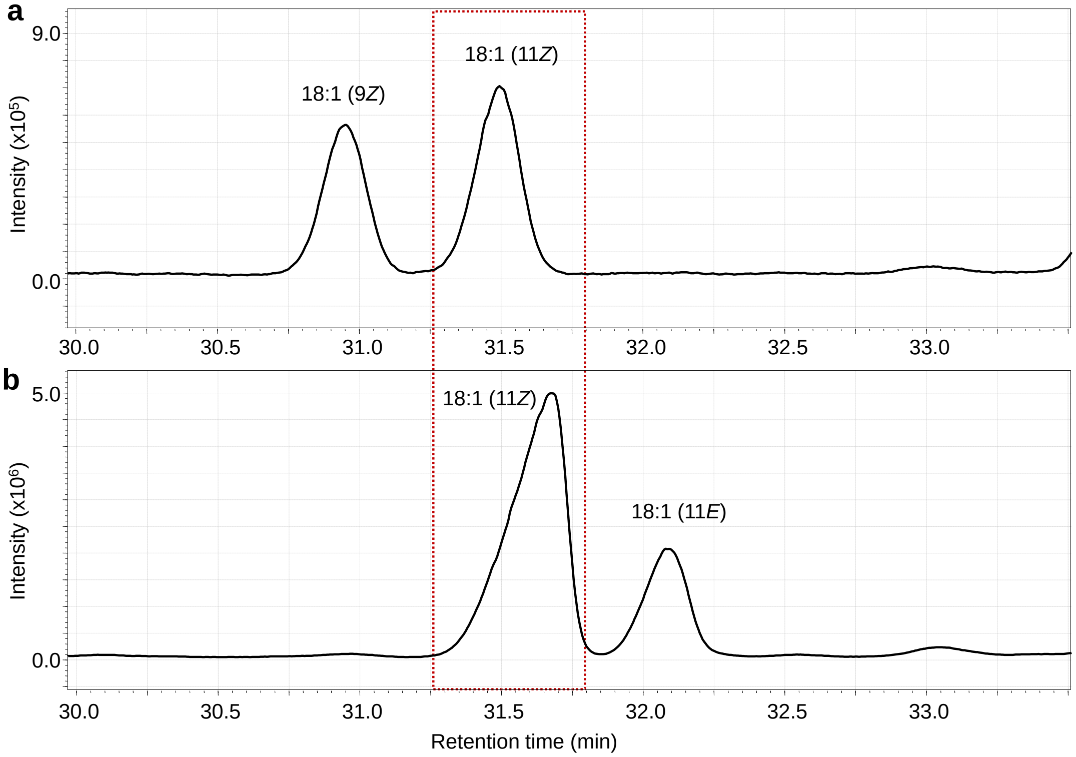


**Figure S17.** GC-MS total ion chromatograms (TIC) of the isomers of FA 18:1. (a) TIC of FA 18:1 (9*Z*) and FA 18:1 (11*Z*). (b) TIC of FA 18:1 (11*Z*) and FA 18:1 (11*E*).

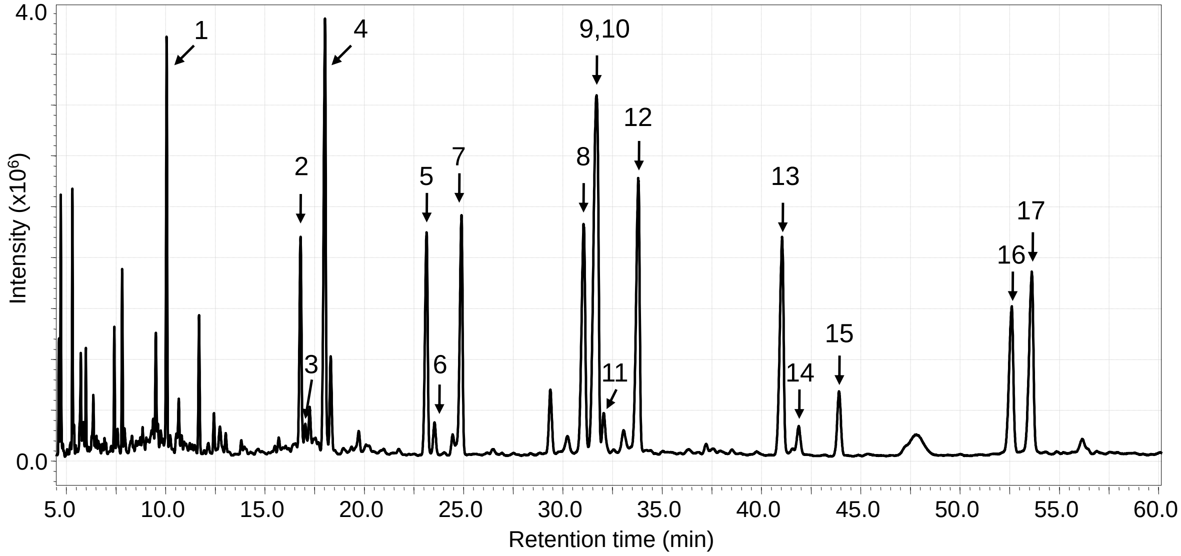


**Figure S18.** GC-MS total ion chromatogram for the analysis of methylated fatty acids standards. **1**: FA 14:0; **2**: FA 16:1 (9*Z*); **3**: FA 16:1 (9*E*); **4**: FA 16:0; **5**: FA 17:1 (10*Z*); **6**: FA 17:1 (10*E*); **7**: FA 17:0; **8**: FA 18:1 (9*Z*); **9**/**10**: FA 18:1 (9*E*)/FA 18:1 (11*Z*); **11**: FA 18:1 (11*E*); **12**: FA 18:0; **13**: FA 19:1 (10*Z*); **14**: FA 19:1 (10*E*); **15**: FA 19:0; **16**: FA 20:1 (11*Z*); **17**: FA 20:1 (11*E*).

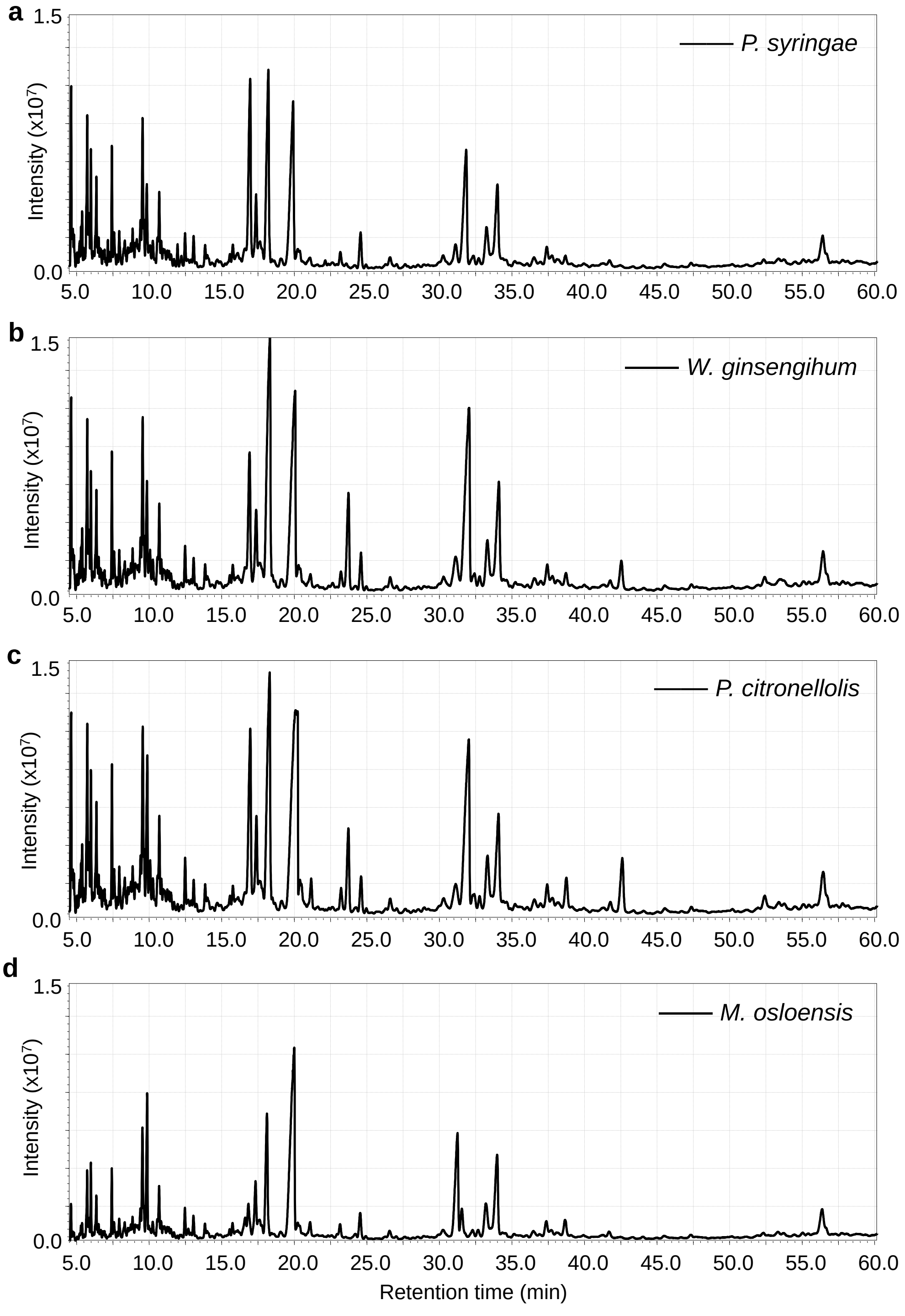


**Figure S19.** GC-MS total ion chromatogram for the analysis of methylated free fatty acids in four bacterial samples. (a) *P. syringae*. (b) *W. ginsengihum*. (c) *P. citronellolis*. (d) *M. osloensis*.

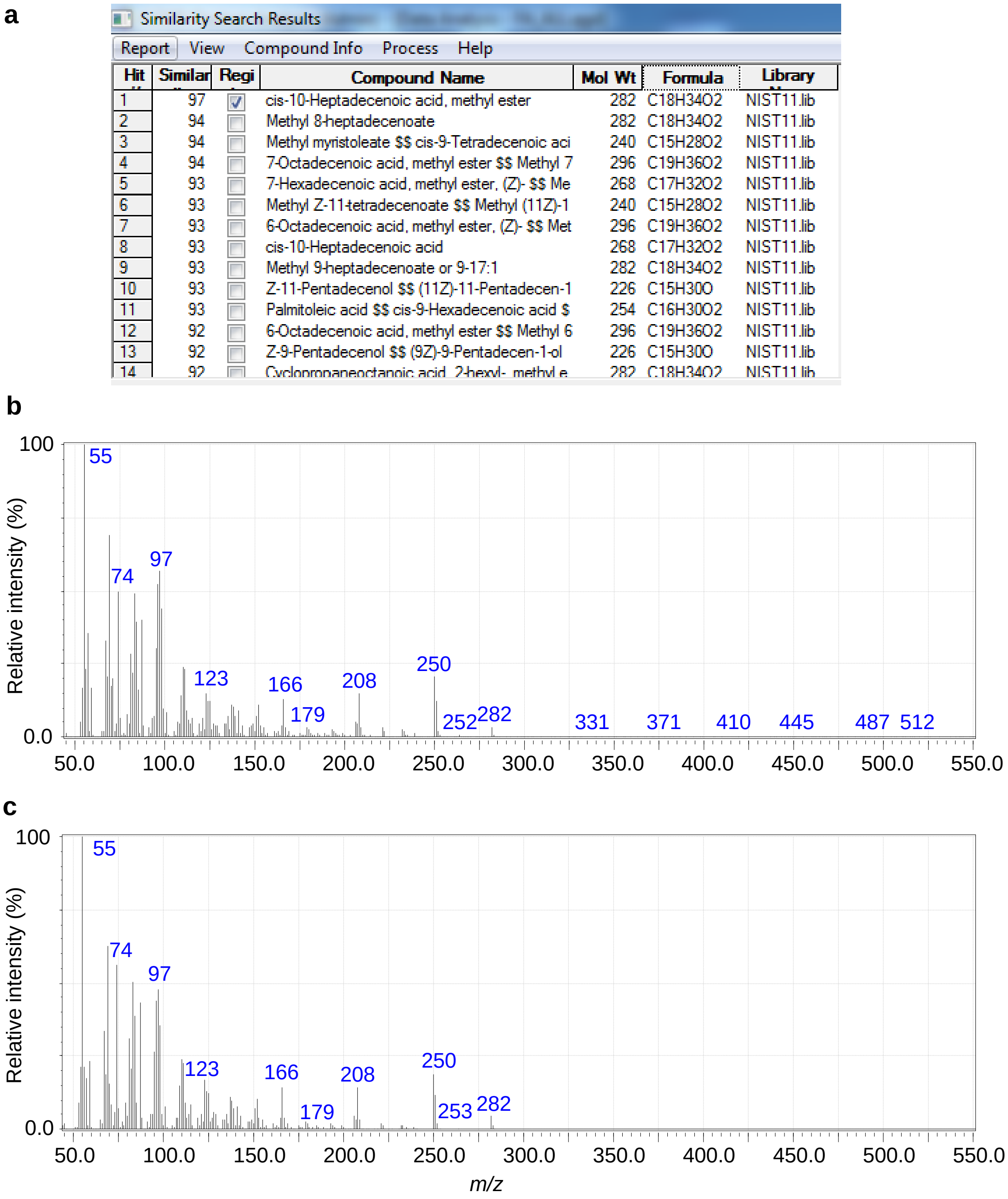

| Hit | Similar | Regi | Compound Name | Mol Wt | Formula | Library |
|---|---|---|---|---|---|---|
| 1 | 97 | ☑ | cis-10-Heptadecenoic acid, methyl ester | 282 | C18H34O2 | NIST11.lib |
| 2 | 94 | ☐ | Methyl 8-heptadecenoate | 282 | C18H34O2 | NIST11.lib |
| 3 | 94 | ☐ | Methyl myristoleate $$ cis-9-Tetradecenoic aci | 240 | C15H28O2 | NIST11.lib |
| 4 | 94 | ☐ | 7-Octadecenoic acid, methyl ester $$ Methyl 7 | 296 | C19H36O2 | NIST11.lib |
| 5 | 93 | ☐ | 7-Hexadecenoic acid, methyl ester, (Z)- $$ Me | 268 | C17H32O2 | NIST11.lib |
| 6 | 93 | ☐ | Methyl Z-11-tetradecenoate $$ Methyl (11Z)-1 | 240 | C15H28O2 | NIST11.lib |
| 7 | 93 | ☐ | 6-Octadecenoic acid, methyl ester, (Z)- $$ Met | 296 | C19H36O2 | NIST11.lib |
| 8 | 93 | ☐ | cis-10-Heptadecenoic acid | 268 | C17H32O2 | NIST11.lib |
| 9 | 93 | ☐ | Methyl 9-heptadecenoate or 9-17:1 | 282 | C18H34O2 | NIST11.lib |
| 10 | 93 | ☐ | Z-11-Pentadecenol $$ (11Z)-11-Pentadecen-1 | 226 | C15H30O | NIST11.lib |
| 11 | 93 | ☐ | Palmitoleic acid $$ cis-9-Hexadecenoic acid $ | 254 | C16H30O2 | NIST11.lib |
| 12 | 92 | ☐ | 6-Octadecenoic acid, methyl ester $$ Methyl 6 | 296 | C19H36O2 | NIST11.lib |
| 13 | 92 | ☐ | Z-9-Pentadecenol $$ (9Z)-9-Pentadecen-1-ol | 226 | C15H30O | NIST11.lib |
| 14 | 92 | ☐ | Cyclopropaneoctanoic acid, 2-hexyl-, methyl e | 282 | C18H34O2 | NIST11.lib |



**Figure S20.** GC-MS analysis of FA 17:1 (10*Z*) standard. (a) Similarity search results from the NIST library. (b) Measured EI mass spectrum of methylated FA 17:1 (10*Z*) standard. (c) The matched EI mass spectrum of methylated FA 17:1 (10*Z*) in NIST library.

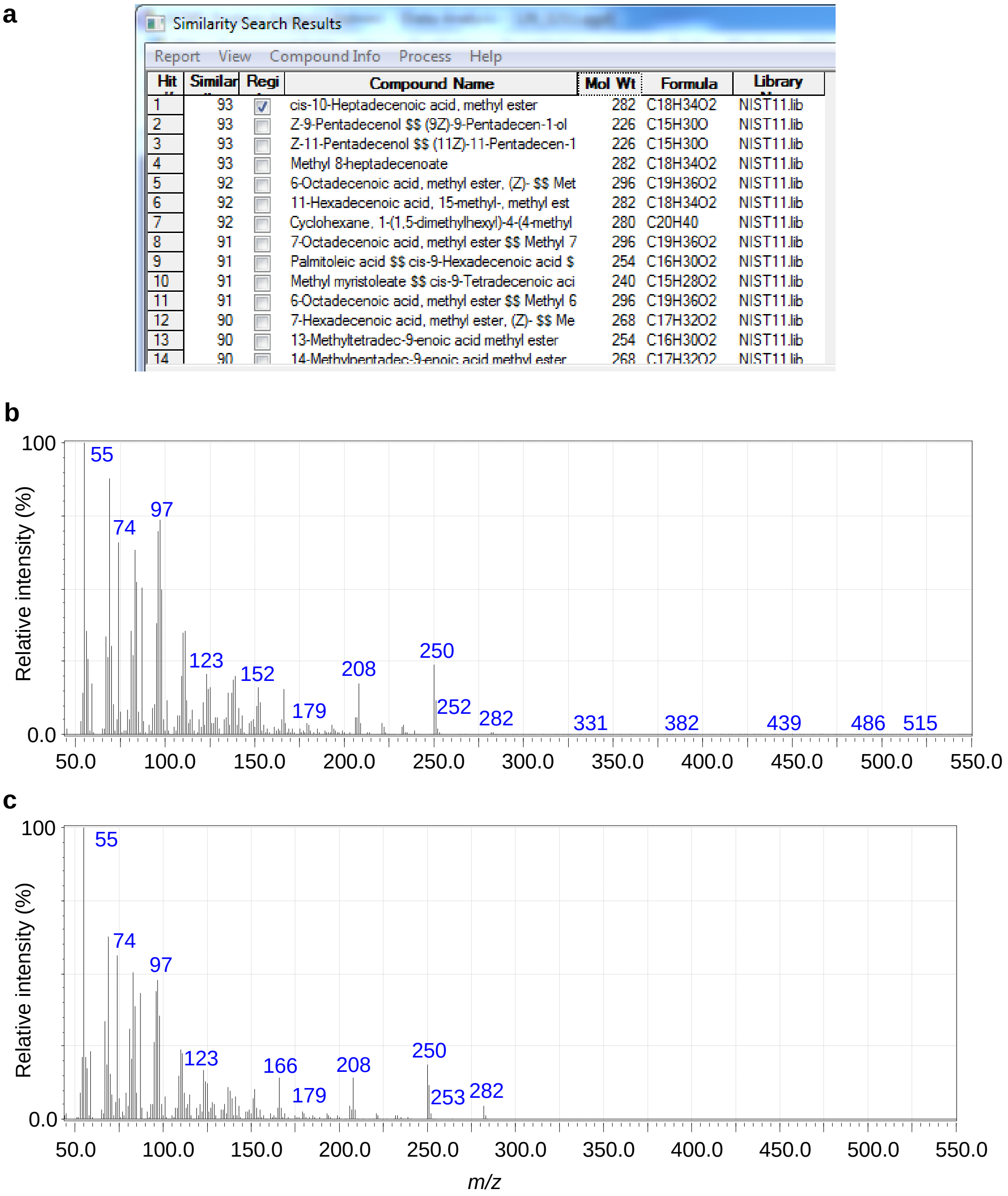

| Hit | Similar | Regi | Compound Name | Mol Wt | Formula | Library |
|---|---|---|---|---|---|---|
| 1 | 93 | ☑ | cis-10-Heptadecenoic acid, methyl ester | 282 | C18H34O2 | NIST11.lib |
| 2 | 93 | ☐ | Z-9-Pentadecenol $$ (9Z)-9-Pentadecen-1-ol | 226 | C15H30O | NIST11.lib |
| 3 | 93 | ☐ | Z-11-Pentadecenol $$ (11Z)-11-Pentadecen-1 | 226 | C15H30O | NIST11.lib |
| 4 | 93 | ☐ | Methyl 8-heptadecenoate | 282 | C18H34O2 | NIST11.lib |
| 5 | 92 | ☐ | 6-Octadecenoic acid, methyl ester, (Z)- $$ Met | 296 | C19H36O2 | NIST11.lib |
| 6 | 92 | ☐ | 11-Hexadecenoic acid, 15-methyl-, methyl est | 282 | C18H34O2 | NIST11.lib |
| 7 | 92 | ☐ | Cyclohexane, 1-(1,5-dimethylhexyl)-4-(4-methyl | 280 | C20H40 | NIST11.lib |
| 8 | 91 | ☐ | 7-Octadecenoic acid, methyl ester $$ Methyl 7 | 296 | C19H36O2 | NIST11.lib |
| 9 | 91 | ☐ | Palmitoleic acid $$ cis-9-Hexadecenoic acid $ | 254 | C16H30O2 | NIST11.lib |
| 10 | 91 | ☐ | Methyl myristoleate $$ cis-9-Tetradecenoic aci | 240 | C15H28O2 | NIST11.lib |
| 11 | 91 | ☐ | 6-Octadecenoic acid, methyl ester $$ Methyl 6 | 296 | C19H36O2 | NIST11.lib |
| 12 | 90 | ☐ | 7-Hexadecenoic acid, methyl ester, (Z)- $$ Me | 268 | C17H32O2 | NIST11.lib |
| 13 | 90 | ☐ | 13-Methyltetradec-9-enoic acid methyl ester | 254 | C16H30O2 | NIST11.lib |
| 14 | 90 | ☐ | 14-Methylpentadec-9-enoic acid methyl ester | 268 | C17H32O2 | NIST11.lib |



**Figure S21.** GC-MS analysis of FA 17:1 (10*Z*) in *W. ginsengihum* bacterial sample. (a) Similarity search results from NIST library. (b) Measured EI mass spectrum of methylated FA 17:1 (10*Z*) in *W. ginsengihum* bacterial sample. (c) The matched EI mass spectrum of methylated FA 17:1 (10*Z*) in NIST library.

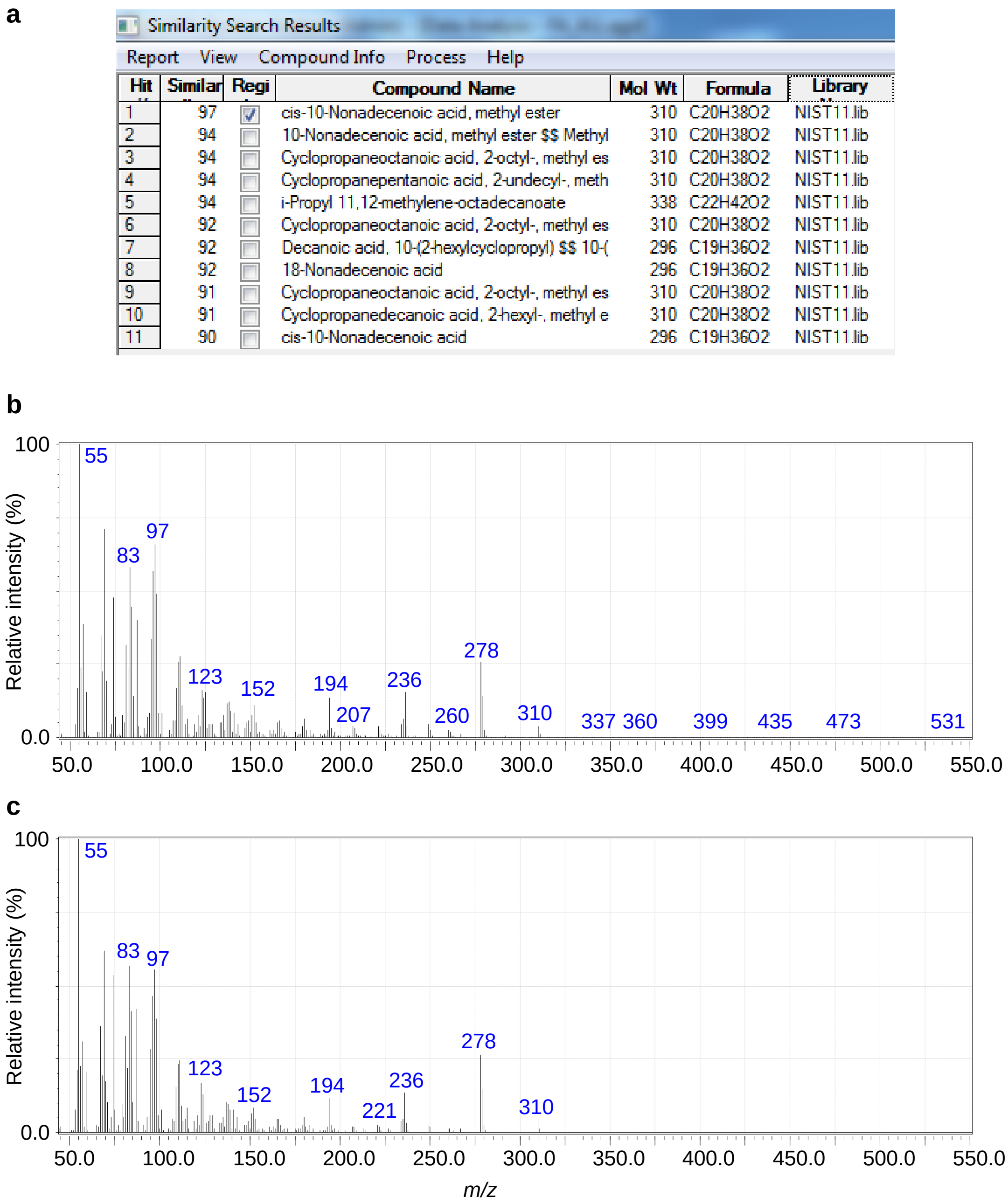

Similarity Search Results

Report View Compound Info Process Help

| Hit | Similar | Regi | Compound Name | Mol Wt | Formula | Library |
|---|---|---|---|---|---|---|
| 1 | 97 | ☑ | cis-10-Nonadecenoic acid, methyl ester | 310 | C20H38O2 | NIST11.lib |
| 2 | 94 | ☐ | 10-Nonadecenoic acid, methyl ester $$ Methyl | 310 | C20H38O2 | NIST11.lib |
| 3 | 94 | ☐ | Cyclopropaneoctanoic acid, 2-octyl-, methyl es | 310 | C20H38O2 | NIST11.lib |
| 4 | 94 | ☐ | Cyclopropanepentanoic acid, 2-undecyl-, meth | 310 | C20H38O2 | NIST11.lib |
| 5 | 94 | ☐ | i-Propyl 11,12-methylene-octadecanoate | 338 | C22H42O2 | NIST11.lib |
| 6 | 92 | ☐ | Cyclopropaneoctanoic acid, 2-octyl-, methyl es | 310 | C20H38O2 | NIST11.lib |
| 7 | 92 | ☐ | Decanoic acid, 10-(2-hexylcyclopropyl) $$ 10-( | 296 | C19H36O2 | NIST11.lib |
| 8 | 92 | ☐ | 18-Nonadecenoic acid | 296 | C19H36O2 | NIST11.lib |
| 9 | 91 | ☐ | Cyclopropaneoctanoic acid, 2-octyl-, methyl es | 310 | C20H38O2 | NIST11.lib |
| 10 | 91 | ☐ | Cyclopropanedecanoic acid, 2-hexyl-, methyl e | 310 | C20H38O2 | NIST11.lib |
| 11 | 90 | ☐ | cis-10-Nonadecenoic acid | 296 | C19H36O2 | NIST11.lib |



**Figure S22.** GC-MS analysis of FA 19:1 (10*Z*) standard. (a) Similarity search results from the NIST library. (b) Measured EI mass spectrum of methylated FA 19:1 (10*Z*) standard. (c) The matched EI mass spectrum of methylated FA 19:1 (10*Z*) in NIST library.

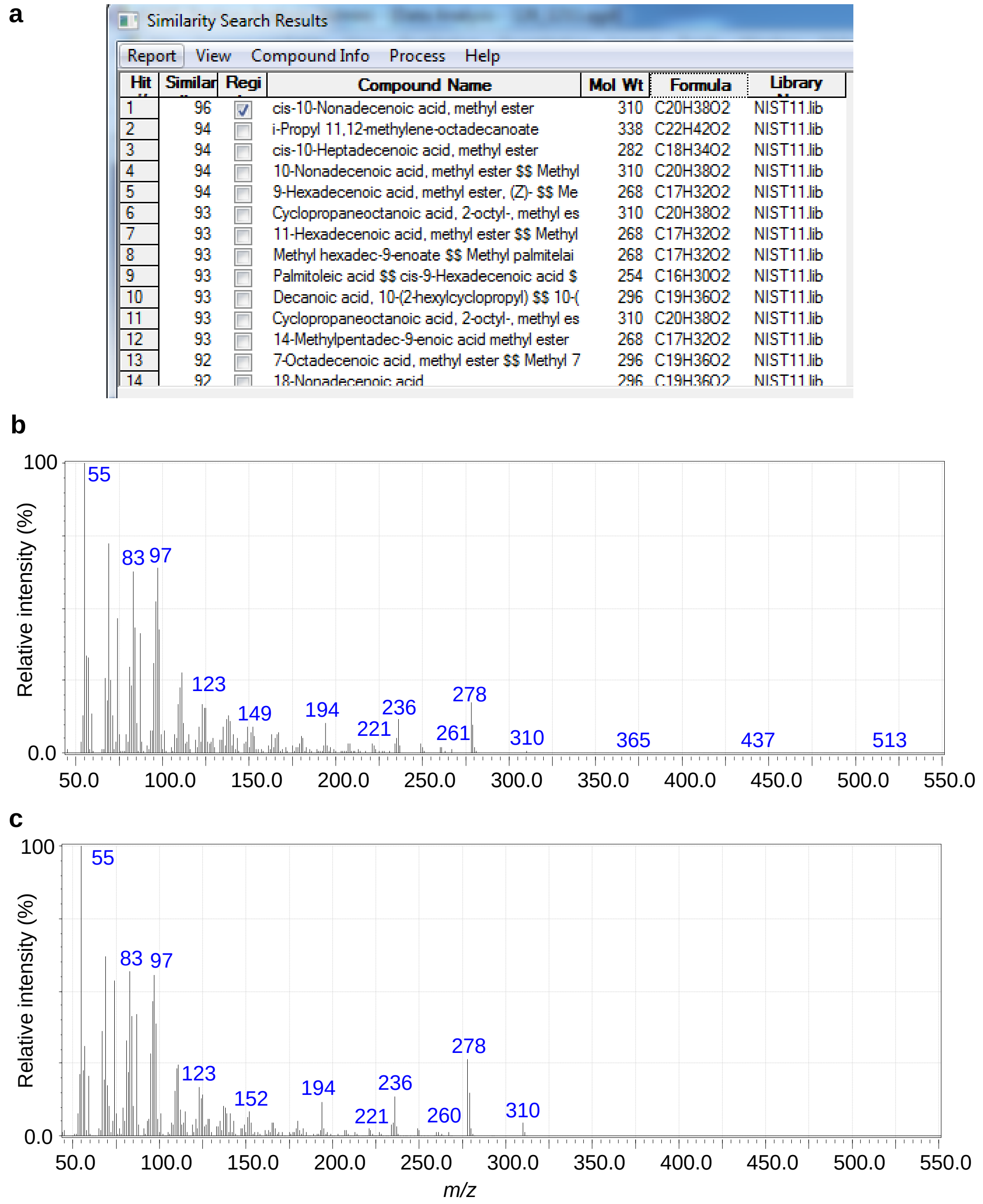

| Hit # | Similar | Regi | Compound Name | Mol Wt | Formula | Library |
|---|---|---|---|---|---|---|
| 1 | 96 | ☑ | cis-10-Nonadecenoic acid, methyl ester | 310 | C20H38O2 | NIST11.lib |
| 2 | 94 | ☐ | i-Propyl 11,12-methylene-octadecanoate | 338 | C22H42O2 | NIST11.lib |
| 3 | 94 | ☐ | cis-10-Heptadecenoic acid, methyl ester | 282 | C18H34O2 | NIST11.lib |
| 4 | 94 | ☐ | 10-Nonadecenoic acid, methyl ester $$ Methyl | 310 | C20H38O2 | NIST11.lib |
| 5 | 94 | ☐ | 9-Hexadecenoic acid, methyl ester, (Z)- $$ Me | 268 | C17H32O2 | NIST11.lib |
| 6 | 93 | ☐ | Cyclopropaneoctanoic acid, 2-octyl-, methyl es | 310 | C20H38O2 | NIST11.lib |
| 7 | 93 | ☐ | 11-Hexadecenoic acid, methyl ester $$ Methyl | 268 | C17H32O2 | NIST11.lib |
| 8 | 93 | ☐ | Methyl hexadec-9-enoate $$ Methyl palmitelai | 268 | C17H32O2 | NIST11.lib |
| 9 | 93 | ☐ | Palmitoleic acid $$ cis-9-Hexadecenoic acid $ | 254 | C16H30O2 | NIST11.lib |
| 10 | 93 | ☐ | Decanoic acid, 10-(2-hexylcyclopropyl) $$ 10-( | 296 | C19H36O2 | NIST11.lib |
| 11 | 93 | ☐ | Cyclopropaneoctanoic acid, 2-octyl-, methyl es | 310 | C20H38O2 | NIST11.lib |
| 12 | 93 | ☐ | 14-Methylpentadec-9-enoic acid methyl ester | 268 | C17H32O2 | NIST11.lib |
| 13 | 92 | ☐ | 7-Octadecenoic acid, methyl ester $$ Methyl 7 | 296 | C19H36O2 | NIST11.lib |
| 14 | 92 | ☐ | 18-Nonadecenoic acid | 296 | C19H36O2 | NIST11.lib |



**Figure S23.** GC-MS analysis of FA 19:1 (10*Z*) in *W. ginsengihum* bacterial sample. (a) Similarity search results from NIST library. (b) Measured EI mass spectrum of methylated FA 19:1 (10*Z*) in *W. ginsengihum* bacterial sample. (c) The matched EI mass spectrum of methylated FA 19:1 (10*Z*) in NIST library.

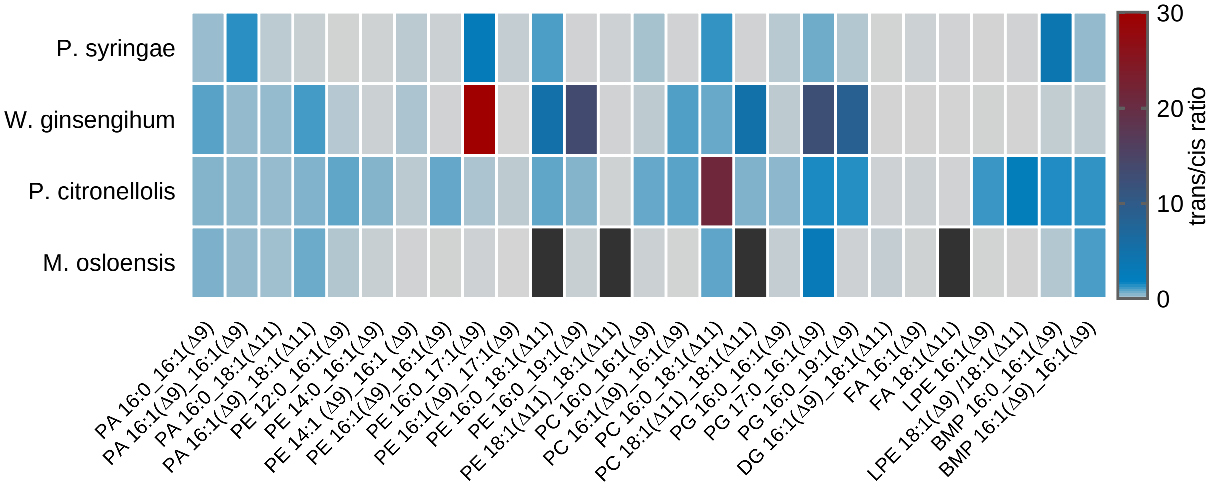


**Figure S24.** Heat map that shows the EIC peak area ratios of lipids *trans*-isomers to their *cis*-counterpart from the four bacterial samples. The black boxes denote the missing data.

**Table S1.** The analytical results of fatty acids in bacterial samples using GC-MS and LC-MS.

| Fatty acids | GC-MS (Peak areas of methylated free FAs in TICs) | | | | LC-MS (Peak areas of FAs in EICs) | | | |
|---|---|---|---|---|---|---|---|---|
| Bacteria | P. syringae | W. ginsengihum | P. citronellolis | M. osloensis | P. syringae | W. ginsengihum | P. citronellolis | M. osloensis |
| 14:0 | | | | | 3.3E+03 | 8.7E+03 | 9.7E+04 | 8.5E+03 |
| 16:0 | 1.5E+08 | 2.8E+08 | 2.5E+08 | 8.2E+07 | 4.9E+05 | 8.6E+05 | 5.7E+05 | 8.9E+05 |
| 16:1 (9*Z*) | 1.4E+08 | 9.8E+07 | 1.5E+08 | 2.4E+07 | 1.6E+05 | 7.1E+05 | 1.8E+06 | 1.0E+06 |
| 16:1 (9*E*) | | | | | 1.1E+04 | 5.4E+03 | 1.4E+05 | 4.2E+04 |
| 17:0 | | | | | 3.5E+03 | 4.7E+03 | 5.1E+04 | 1.9E+04 |
| 17:1 (10*Z*) | 1.2E+07 | 1.5E+07 | 1.9E+07 | 1.1E+07 | 2.2E+03 | 7.9E+02 | 8.5E+04 | 1.6E+05 |
| 17:1 (10*E*) | | 7.2E+07 | 6.0E+07 | | 1.2E+04 | 3.5E+04 | 2.7E+04 | 6.1E+03 |
| 18:0 | 9.5E+07 | 1.3E+08 | 1.2E+08 | 8.9E+07 | 3.1E+05 | 7.7E+05 | 1.0E+06 | 1.0E+06 |
| 18:1 (9*Z*) | | | | 1.0E+08 | | | | 2.6E+06 |
| 18:1 (9*E*) | | | | 2.5E+07 | | | | 1.4E+04 |
| 18:1 (11*Z*) | 1.3E+08 | 2.7E+08 | 2.5E+08 | | 1.7E+05 | 8.8E+05 | 4.0E+06 | |
| 18:1 (11*E*) | 1.2E+07 | 1.8E+07 | 2.3E+07 | | 4.1E+03 | 5.0E+03 | 1.4E+05 | |
| 19:0 | | | | | 1.2E+03 | 1.3E+04 | 9.4E+04 | 6.5E+02 |
| 19:1 (10*Z*) | 6.9E+06 | 1.1E+07 | 1.4E+07 | 6.4E+06 | 3.4E+03 | 1.5E+02 | 8.9E+03 | 7.0E+03 |
| 19:1 (10*E*) | | 2.8E+07 | 5.7E+07 | | | 1.3E+04 | 6.9E+04 | |
| 20:0 | | | | | 5.6E+03 | 2.3E+04 | 2.0E+04 | 1.6E+04 |
| 20:1 (11*Z*) | | | | | 4.0E+03 | 4.3E+03 | 4.3E+04 | 6.4E+03 |
| 20:1 (11*E*) | | | | | | | 3.9E+03 | |

**Table S2**. EIC peak area ratios of lipids *trans*-isomers to their *cis*-counterpart from the four bacterial samples

| Lipids | P. syringae | W. ginsengihum | P. citronellolis | M. osloensis |
|---|---|---|---|---|
| PA 16:0_16:1(Δ9) | 0.55 | 1.17 | 0.76 | 0.82 |
| PA 16:1(Δ9)_16:1(Δ9) | 1.60 | 0.61 | 0.63 | 0.61 |
| PA 16:0_18:1(Δ11) | 0.22 | 0.59 | 0.58 | 0.48 |
| PA 16:1(Δ9)_18:1(Δ11) | 0.11 | 1.36 | 0.77 | 0.97 |
| PE 12:0_16:1(Δ9) | 0.02 | 0.28 | 1.09 | 0.31 |
| PE 14:0_16:1(Δ9) | 0.04 | 0.08 | 0.76 | 0.11 |
| PE 14:1 (Δ9)_16:1 (Δ9) | 0.23 | 0.37 | 0.23 | 0.03 |
| PE 16:1(Δ9)_16:1(Δ9) | 0.08 | 0.03 | 1.05 | 0.02 |
| PE 16:0_17:1(Δ9) | 2.98 | 30.05 | 0.37 | 0.06 |
| PE 16:1(Δ9)_17:1(Δ9) | 0.15 | 0.00 | 0.21 | 0.00 |
| PE 16:0_18:1(Δ11) | 1.27 | 5.40 | 1.09 | N/A |
| PE 16:0_19:1(Δ9) | 0.03 | 13.74 | 0.75 | 0.12 |
| PE 18:1(Δ11)_18:1(Δ11)[1] | 0.06 | 0.03 | 0.05 | N/A |
| PC 16:0_16:1(Δ9) | 0.43 | 0.21 | 1.04 | 0.08 |
| PC 16:1(Δ9)_16:1(Δ9) | 0.03 | 1.25 | 1.15 | 0.01 |
| PC 16:0_18:1(Δ11) | 1.51 | 1.01 | 21.32 | 1.10 |
| PC 18:1(Δ11)_18:1(Δ11)[2] | 0.03 | 5.25 | 0.79 | N/A |
| PG 16:0_16:1(Δ9) | 0.26 | 0.21 | 0.67 | 0.15 |
| PG 17:0_16:1(Δ9) | 0.95 | 12.71 | 1.74 | 3.17 |
| PG 16:0_19:1(Δ9) | 0.30 | 8.85 | 1.64 | 0.06 |
| DG 16:1(Δ9)_18:1(Δ11) | 0.02 | 0.01 | 0.06 | 0.16 |
| FA 16:1(Δ9) | 0.07 | 0.01 | 0.08 | 0.04 |
| FA 18:1(Δ11) | 0.02 | 0.01 | 0.03 | N/A |
| LPE 16:1(Δ9) | 0.01 | 0.02 | 1.46 | 0.01 |
| LPE 18:1(Δ9) /18:1(Δ11) | 0.03 | 0.01 | 2.63 | 0.01 |
| BMP 16:0_16:1(Δ9) | 4.09 | 0.16 | 1.69 | 0.31 |
| BMP 16:1(Δ9)_16:1(Δ9) | 0.60 | 0.21 | 1.53 | 1.30 |

Note: [1]ratio of PE 18:1 (11*Z*)_18:1 (11*E*) to PE 18:1 (11*Z*)_18:1 (11*Z*); [2] ratio of PC 18:1 (11*Z*)_18:1 (11*E*) to PC 18:1 (11*Z*)_18:1 (11*Z*). N/A means not detected.

**Appendix: Cartesian coordinates and energies of optimized structures**

**1**

M06-2X SCF energy: -856.43105458 a.u.

M06-2X enthalpy: -855.903711 a.u.

M06-2X free energy: -855.982342 a.u.

M06-2X SCF energy in solution: -856.71684890 a.u.

M06-2X enthalpy in solution: -856.189505 a.u.

M06-2X free energy in solution: -856.268136 a.u.

Cartesian coordinates

| ATOM | X | Y | Z |
|---|---|---|---|
| C | -0.620363 | 4.163026 | 0.453516 |
| C | -1.939955 | 3.963518 | 0.463309 |
| H | -0.208787 | 4.861929 | 1.182193 |
| C | -2.729939 | 3.021145 | -0.399940 |
| C | -3.300804 | 1.848906 | 0.408983 |
| H | -2.119096 | 2.628225 | -1.219610 |
| H | -3.563209 | 3.565709 | -0.865097 |
| C | -4.166669 | 0.913349 | -0.431379 |
| H | -2.469745 | 1.286579 | 0.854820 |
| H | -3.892349 | 2.239451 | 1.248369 |
| H | -4.994934 | 1.484391 | -0.873872 |
| H | -3.575274 | 0.528705 | -1.274165 |
| C | -4.731026 | -0.259973 | 0.366957 |
| C | -5.599328 | -1.195115 | -0.472141 |
| H | -5.321137 | 0.124797 | 1.210400 |
| H | -3.902529 | -0.831251 | 0.808305 |
| C | -6.163482 | -2.369041 | 0.325120 |
| H | -6.428061 | -0.623473 | -0.912905 |
| H | -5.009373 | -1.579134 | -1.316167 |
| H | -5.335173 | -2.941926 | 0.765307 |
| H | -6.752916 | -1.985737 | 1.169946 |
| C | -7.033275 | -3.303725 | -0.513325 |
| C | -7.589636 | -4.472184 | 0.296365 |
| H | -6.443189 | -3.685254 | -1.356556 |
| H | -7.860008 | -2.729932 | -0.951554 |
| H | -6.778610 | -5.073929 | 0.720109 |
| H | -8.204170 | -4.112262 | 1.128384 |
| H | -8.209836 | -5.130320 | -0.318902 |
| C | 0.399450 | 3.493407 | -0.423776 |

| | | | |
|---|---|---|---|
| C | 1.275005 | 2.512337 | 0.366853 |
| H | 1.046848 | 4.256789 | -0.876717 |
| H | -0.079239 | 2.960660 | -1.252199 |
| C | 2.369302 | 1.872973 | -0.484644 |
| H | 1.732404 | 3.035887 | 1.217469 |
| H | 0.633795 | 1.731337 | 0.796333 |
| H | 1.910710 | 1.357209 | -1.339996 |
| H | 3.007439 | 2.661202 | -0.908189 |
| C | 3.234164 | 0.886245 | 0.296646 |
| C | 4.331918 | 0.248220 | -0.551997 |
| H | 2.596345 | 0.097031 | 0.717772 |
| H | 3.690803 | 1.400757 | 1.153310 |
| C | 5.188672 | -0.739337 | 0.235974 |
| H | 3.875655 | -0.264385 | -1.410523 |
| H | 4.971785 | 1.037359 | -0.971280 |
| H | 5.649623 | -0.236101 | 1.093815 |
| H | 4.558518 | -1.532226 | 0.655275 |
| C | 6.280239 | -1.369317 | -0.618939 |
| H | 6.949943 | -0.608923 | -1.039451 |
| H | 5.858148 | -1.902114 | -1.480060 |
| C | 7.126215 | -2.347925 | 0.156279 |
| O | 6.991120 | -2.630764 | 1.319849 |
| O | 8.087693 | -2.897088 | -0.614318 |
| H | 8.581143 | -3.509473 | -0.040849 |
| H | -2.528953 | 4.512193 | 1.198866 |

**2**

M06-2X SCF energy: -573.08590630 a.u.
M06-2X enthalpy: -572.921763 a.u.
M06-2X free energy: -572.970627 a.u.
M06-2X SCF energy in solution: -573.27062379 a.u.
M06-2X enthalpy in solution: -573.106480 a.u.
M06-2X free energy in solution: -573.155345 a.u.

Cartesian coordinates

| ATOM | X | Y | Z |
|---|---|---|---|
| C | 2.918128 | 1.187930 | -0.193238 |
| C | 1.537056 | 1.291691 | -0.128114 |
| C | 0.742146 | 0.141413 | 0.051605 |
| C | 1.367823 | -1.111058 | 0.183034 |
| C | 2.752161 | -1.196379 | 0.106405 |

| | | | |
|---|---|---|---|
| C | 3.533782 | -0.057349 | -0.079478 |
| H | 3.515548 | 2.081960 | -0.340530 |
| H | 1.059359 | 2.259961 | -0.237075 |
| H | 0.770515 | -1.998351 | 0.336943 |
| H | 3.225289 | -2.168595 | 0.201132 |
| H | 4.614447 | -0.139490 | -0.133933 |
| C | -0.709601 | 0.319927 | 0.127956 |
| C | -1.690319 | -0.777330 | -0.040015 |
| O | -1.114528 | 1.533792 | 0.374141 |
| O | -1.334120 | -1.926000 | -0.174727 |
| C | -3.525348 | 0.815410 | -0.144857 |
| H | -3.099734 | 1.334681 | -1.008395 |
| H | -3.361063 | 1.399244 | 0.765215 |
| H | -4.597948 | 0.687732 | -0.296461 |
| O | -3.011525 | -0.500625 | -0.022749 |

**3**

M06-2X SCF energy: -1429.57321516 a.u.

M06-2X enthalpy: -1428.878323 a.u.

M06-2X free energy: -1428.980637 a.u.

M06-2X SCF energy in solution: -1430.03546970 a.u.

M06-2X enthalpy in solution: -1429.340578 a.u.

M06-2X free energy in solution: -1429.442892 a.u.

Cartesian coordinates

| ATOM | X | Y | Z |
|---|---|---|---|
| C | 0.102157 | -0.661432 | -0.594688 |
| C | 0.604314 | 0.526130 | -1.343699 |
| H | -0.140510 | -1.473141 | -1.293045 |
| H | 0.419525 | 0.592146 | -2.410549 |
| C | 1.431594 | 1.568110 | -0.668367 |
| C | 2.936366 | 1.392988 | -0.945204 |
| H | 1.271773 | 1.524765 | 0.416297 |
| H | 1.128076 | 2.570275 | -1.001678 |
| C | 3.786670 | 2.472743 | -0.280351 |
| H | 3.241271 | 0.402259 | -0.586530 |
| H | 3.107619 | 1.404971 | -2.030349 |
| H | 3.459862 | 3.462329 | -0.629951 |
| H | 3.610616 | 2.455526 | 0.804089 |
| C | 5.280293 | 2.308020 | -0.553213 |
| C | 6.134829 | 3.383006 | 0.115262 |

| | | | |
|---|---|---|---|
| H | 5.455831 | 2.326830 | -1.637958 |
| H | 5.605298 | 1.317234 | -0.207034 |
| C | 7.627959 | 3.222349 | -0.161694 |
| H | 5.806252 | 4.374177 | -0.227667 |
| H | 5.962060 | 3.361505 | 1.200353 |
| H | 7.957018 | 2.230838 | 0.179723 |
| H | 7.801486 | 3.245149 | -1.246833 |
| C | 8.483594 | 4.295845 | 0.508169 |
| C | 9.972984 | 4.123887 | 0.220800 |
| H | 8.310262 | 4.270411 | 1.591709 |
| H | 8.152491 | 5.285270 | 0.167218 |
| H | 10.329103 | 3.151720 | 0.577855 |
| H | 10.170653 | 4.173876 | -0.855242 |
| H | 10.570168 | 4.899851 | 0.708245 |
| C | -1.118733 | -0.372048 | 0.284196 |
| C | -2.331592 | 0.077622 | -0.526370 |
| H | -0.841036 | 0.393053 | 1.019706 |
| H | -1.358849 | -1.280656 | 0.852015 |
| C | -3.567773 | 0.303826 | 0.341717 |
| H | -2.088241 | 1.004398 | -1.063090 |
| H | -2.561143 | -0.673722 | -1.295464 |
| H | -3.815788 | -0.626106 | 0.871733 |
| H | -3.334721 | 1.046663 | 1.117088 |
| C | -4.781466 | 0.770426 | -0.458904 |
| C | -6.018054 | 0.998719 | 0.407589 |
| H | -5.014066 | 0.028106 | -1.234798 |
| H | -4.532853 | 1.700347 | -0.988581 |
| C | -7.225878 | 1.468421 | -0.399091 |
| H | -6.268152 | 0.068195 | 0.936226 |
| H | -5.784668 | 1.739586 | 1.185143 |
| H | -6.985944 | 2.398049 | -0.928065 |
| H | -7.467909 | 0.733886 | -1.175912 |
| C | -8.452847 | 1.693595 | 0.474398 |
| H | -8.258722 | 2.440296 | 1.254341 |
| H | -8.741373 | 0.777834 | 1.005133 |
| C | -9.647178 | 2.159709 | -0.319808 |
| O | -9.673675 | 2.347318 | -1.509879 |
| O | -10.725363 | 2.353477 | 0.467174 |
| H | -11.442711 | 2.650756 | -0.119708 |
| O | 1.180418 | -1.103853 | 0.273194 |
| C | 1.221433 | -2.411237 | 0.600422 |

| | | | |
|---|---|---|---|
| C | 1.559725 | -3.380729 | -0.414810 |
| C | 1.018780 | -2.733569 | 2.016133 |
| C | 2.050078 | -2.910669 | -1.654396 |
| C | 1.422873 | -4.774247 | -0.231425 |
| O | 1.125737 | -3.840919 | 2.510040 |
| O | 0.681754 | -1.645627 | 2.738662 |
| C | 2.384208 | -3.796204 | -2.666716 |
| H | 2.185212 | -1.842003 | -1.796841 |
| C | 1.760253 | -5.648172 | -1.254829 |
| H | 1.057082 | -5.151309 | 0.714105 |
| C | 0.480516 | -1.905359 | 4.122996 |
| C | 2.238873 | -5.170398 | -2.474778 |
| H | 2.767560 | -3.415316 | -3.608182 |
| H | 1.647114 | -6.716501 | -1.099163 |
| H | 1.389251 | -2.310854 | 4.574102 |
| H | -0.330406 | -2.623584 | 4.266800 |
| H | 0.226274 | -0.945127 | 4.569496 |
| H | 2.500410 | -5.863128 | -3.268297 |

**3a**

M06-2X SCF energy: -1429.57193648 a.u.

M06-2X enthalpy: -1428.876686 a.u.

M06-2X free energy: -1428.977493 a.u.

M06-2X SCF energy in solution: -1430.03421095 a.u.

M06-2X enthalpy in solution: -1429.338960 a.u.

M06-2X free energy in solution: -1429.439768 a.u.

Cartesian coordinates

| ATOM | X | Y | Z |
|---|---|---|---|
| C | 0.858858 | -2.111297 | -1.328140 |
| C | 1.589585 | -2.480246 | -0.077662 |
| H | 0.535775 | -2.930915 | -1.962171 |
| C | 2.310096 | -1.323116 | 0.605442 |
| C | 3.560540 | -0.901464 | -0.167055 |
| H | 2.582300 | -1.648662 | 1.614898 |
| H | 1.628471 | -0.473481 | 0.729106 |
| C | 4.236921 | 0.329240 | 0.432763 |
| H | 4.273067 | -1.737961 | -0.182028 |
| H | 3.307613 | -0.701209 | -1.217560 |
| H | 3.530452 | 1.171260 | 0.423130 |
| H | 4.470056 | 0.136709 | 1.489143 |

| | | | |
|---|---|---|---|
| C | 5.512919 | 0.730090 | -0.303897 |
| C | 6.189910 | 1.962282 | 0.292768 |
| H | 5.278416 | 0.919700 | -1.360730 |
| H | 6.218106 | -0.112622 | -0.293129 |
| C | 7.467491 | 2.360511 | -0.442671 |
| H | 5.485671 | 2.805944 | 0.281249 |
| H | 6.423090 | 1.772404 | 1.349800 |
| H | 8.171211 | 1.516282 | -0.432840 |
| H | 7.234710 | 2.552081 | -1.499650 |
| C | 8.148039 | 3.590753 | 0.154591 |
| C | 9.423306 | 3.975957 | -0.590764 |
| H | 8.379899 | 3.397365 | 1.209849 |
| H | 7.444115 | 4.432874 | 0.144021 |
| H | 10.150161 | 3.157082 | -0.568507 |
| H | 9.208531 | 4.199464 | -1.641157 |
| H | 9.897348 | 4.857621 | -0.150056 |
| C | 0.394411 | -0.731134 | -1.675877 |
| C | -0.842583 | -0.261730 | -0.885097 |
| H | 0.155373 | -0.700859 | -2.745571 |
| H | 1.196378 | 0.003856 | -1.516182 |
| C | -1.409427 | 1.057777 | -1.401023 |
| H | -1.619833 | -1.036095 | -0.921228 |
| H | -0.567253 | -0.145939 | 0.171500 |
| H | -0.614876 | 1.817174 | -1.424757 |
| H | -1.743494 | 0.928947 | -2.439947 |
| C | -2.572051 | 1.563905 | -0.549645 |
| C | -3.201323 | 2.844240 | -1.094390 |
| H | -2.219581 | 1.739699 | 0.476317 |
| H | -3.338111 | 0.780374 | -0.475099 |
| C | -4.340154 | 3.361919 | -0.219635 |
| H | -2.427863 | 3.619877 | -1.187018 |
| H | -3.573584 | 2.660359 | -2.112020 |
| H | -5.111626 | 2.590686 | -0.112155 |
| H | -3.974104 | 3.564073 | 0.793509 |
| C | -4.972185 | 4.627343 | -0.784212 |
| H | -5.383829 | 4.458244 | -1.786804 |
| H | -4.231058 | 5.428369 | -0.898862 |
| C | -6.084042 | 5.155397 | 0.087059 |
| O | -6.447500 | 4.680729 | 1.133214 |
| O | -6.652351 | 6.258593 | -0.442494 |
| H | -7.350303 | 6.531188 | 0.178728 |

| | | | |
|---|---|---|---|
| O | 0.746662 | -3.176697 | 0.910815 |
| C | -0.592384 | -3.022549 | 0.900909 |
| C | -1.385945 | -3.936279 | 0.113714 |
| C | -1.158382 | -1.994627 | 1.782718 |
| C | -0.721895 | -4.999746 | -0.539067 |
| C | -2.784710 | -3.819494 | -0.051839 |
| O | -2.340986 | -1.743117 | 1.920406 |
| O | -0.195347 | -1.306818 | 2.432133 |
| C | -1.420631 | -5.899039 | -1.327607 |
| H | 0.347833 | -5.110520 | -0.396735 |
| C | -3.470747 | -4.726005 | -0.848331 |
| H | -3.313347 | -3.018451 | 0.446636 |
| C | -0.684208 | -0.264485 | 3.268010 |
| C | -2.800522 | -5.766426 | -1.490601 |
| H | -0.890564 | -6.711793 | -1.814423 |
| H | -4.544304 | -4.618633 | -0.968721 |
| H | -1.340280 | -0.667790 | 4.042645 |
| H | -1.246947 | 0.466325 | 2.680333 |
| H | 0.197482 | 0.196019 | 3.711694 |
| H | -3.347775 | -6.470358 | -2.109453 |
| H | 2.333508 | -3.250871 | -0.321100 |

**4**

M06-2X SCF energy: -1429.57254044 a.u.

M06-2X enthalpy: -1428.877521 a.u.

M06-2X free energy: -1428.979532 a.u.

M06-2X SCF energy in solution: -1430.03593103 a.u.

M06-2X enthalpy in solution: -1429.340912 a.u.

M06-2X free energy in solution: -1429.442923 a.u.

Cartesian coordinates

| ATOM | X | Y | Z |
|---|---|---|---|
| C | -2.122196 | -0.249729 | -0.890359 |
| C | -1.690110 | 0.788419 | -1.865023 |
| H | -2.139685 | 0.179690 | 0.121272 |
| H | -1.556286 | 0.474786 | -2.897271 |
| C | -1.374423 | 2.194343 | -1.475998 |
| C | 0.126056 | 2.434925 | -1.220125 |
| H | -1.927797 | 2.462638 | -0.565119 |
| H | -1.703929 | 2.885325 | -2.263684 |
| C | 0.437376 | 3.890736 | -0.881195 |

| | | | |
|---|---|---|---|
| H | 0.459420 | 1.785241 | -0.399926 |
| H | 0.696539 | 2.130406 | -2.107960 |
| H | 0.097822 | 4.535960 | -1.703443 |
| H | -0.142218 | 4.189664 | 0.003341 |
| C | 1.921786 | 4.139616 | -0.622249 |
| C | 2.235080 | 5.595209 | -0.281941 |
| H | 2.500116 | 3.840595 | -1.507478 |
| H | 2.261513 | 3.493035 | 0.198670 |
| C | 3.719866 | 5.846588 | -0.028679 |
| H | 1.891483 | 6.242047 | -1.101316 |
| H | 1.659246 | 5.892932 | 0.605489 |
| H | 4.064798 | 5.198847 | 0.789488 |
| H | 4.295893 | 5.550386 | -0.916626 |
| C | 4.033917 | 7.301710 | 0.313653 |
| C | 5.521516 | 7.538524 | 0.561270 |
| H | 3.459342 | 7.594992 | 1.201684 |
| H | 3.687111 | 7.947058 | -0.503697 |
| H | 5.881547 | 6.921973 | 1.391690 |
| H | 6.110490 | 7.277393 | -0.324324 |
| H | 5.728199 | 8.584433 | 0.805431 |
| C | -1.265480 | -1.515064 | -0.903194 |
| C | 0.191510 | -1.253708 | -0.530710 |
| H | -1.333620 | -1.962150 | -1.902845 |
| H | -1.710094 | -2.238516 | -0.207744 |
| C | 1.018349 | -2.535830 | -0.453844 |
| H | 0.642827 | -0.573249 | -1.265694 |
| H | 0.237429 | -0.734126 | 0.437711 |
| H | 0.573516 | -3.211818 | 0.289272 |
| H | 0.963570 | -3.060319 | -1.417809 |
| C | 2.480997 | -2.281192 | -0.096900 |
| C | 3.309852 | -3.561510 | -0.020382 |
| H | 2.534904 | -1.755857 | 0.866594 |
| H | 2.925292 | -1.604953 | -0.840146 |
| C | 4.770501 | -3.299227 | 0.337248 |
| H | 2.865176 | -4.238257 | 0.722675 |
| H | 3.255999 | -4.087030 | -0.984153 |
| H | 5.223012 | -2.626074 | -0.400111 |
| H | 4.834224 | -2.778131 | 1.299520 |
| C | 5.587438 | -4.582415 | 0.409320 |
| H | 5.567191 | -5.125141 | -0.543861 |
| H | 5.179803 | -5.276332 | 1.154873 |

| | | | |
|---|---|---|---|
| C | 7.031450 | -4.326946 | 0.761415 |
| O | 7.526422 | -3.248604 | 0.972016 |
| O | 7.741246 | -5.472446 | 0.819104 |
| H | 8.652181 | -5.218880 | 1.049972 |
| O | -3.482559 | -0.648714 | -1.239514 |
| C | -4.298888 | -0.999136 | -0.226449 |
| C | -4.749894 | 0.016727 | 0.697331 |
| C | -4.755394 | -2.391117 | -0.199674 |
| C | -4.556340 | 1.369888 | 0.342226 |
| C | -5.363963 | -0.272695 | 1.934338 |
| O | -5.569935 | -2.848389 | 0.581048 |
| O | -4.157514 | -3.139848 | -1.148810 |
| C | -4.947120 | 2.391225 | 1.193844 |
| H | -4.123832 | 1.596691 | -0.627634 |
| C | -5.749402 | 0.759879 | 2.777888 |
| H | -5.530948 | -1.303866 | 2.215854 |
| C | -4.578521 | -4.498622 | -1.169704 |
| C | -5.541907 | 2.091990 | 2.419972 |
| H | -4.798047 | 3.424985 | 0.897296 |
| H | -6.216192 | 0.522182 | 3.728677 |
| H | -5.654161 | -4.565036 | -1.349507 |
| H | -4.353455 | -4.987093 | -0.218415 |
| H | -4.024482 | -4.966819 | -1.982036 |
| H | -5.847528 | 2.890869 | 3.088060 |

**4a**

M06-2X SCF energy: -1429.57486523 a.u.
M06-2X enthalpy: -1428.879843 a.u.
M06-2X free energy: -1428.981637 a.u.
M06-2X SCF energy in solution: -1430.03807846 a.u.
M06-2X enthalpy in solution: -1429.343056 a.u.
M06-2X free energy in solution: -1429.44485 a.u.

Cartesian coordinates

| ATOM | X | Y | Z |
|---|---|---|---|
| C | -1.723417 | -0.040496 | -1.363551 |
| C | -2.387229 | 0.557055 | -0.171586 |
| H | -1.906324 | 0.422729 | -2.328547 |
| C | -2.359694 | 2.077824 | -0.147542 |
| C | -0.941411 | 2.635246 | -0.051361 |
| H | -2.960098 | 2.421668 | 0.703713 |

| | | | |
|---|---|---|---|
| H | -2.854884 | 2.445621 | -1.055281 |
| C | -0.910869 | 4.160726 | 0.018810 |
| H | -0.444886 | 2.220771 | 0.837969 |
| H | -0.353423 | 2.296540 | -0.915382 |
| H | -1.407773 | 4.574898 | -0.869350 |
| H | -1.498776 | 4.497251 | 0.883898 |
| C | 0.504335 | 4.725669 | 0.117980 |
| C | 0.538028 | 6.250875 | 0.188948 |
| H | 1.092089 | 4.388529 | -0.747146 |
| H | 1.000200 | 4.309459 | 1.006019 |
| C | 1.952712 | 6.816101 | 0.293883 |
| H | 0.045022 | 6.667253 | -0.700624 |
| H | -0.052825 | 6.587668 | 1.052217 |
| H | 2.445754 | 6.400250 | 1.183841 |
| H | 2.544591 | 6.479214 | -0.568786 |
| C | 1.987863 | 8.341544 | 0.364678 |
| C | 3.407855 | 8.891654 | 0.471151 |
| H | 1.395538 | 8.676025 | 1.226087 |
| H | 1.496261 | 8.755070 | -0.525312 |
| H | 3.906801 | 8.511663 | 1.368940 |
| H | 4.008400 | 8.590541 | -0.393741 |
| H | 3.414399 | 9.984264 | 0.520398 |
| C | -0.842796 | -1.241292 | -1.260066 |
| C | 0.629048 | -0.873942 | -0.991261 |
| H | -0.897252 | -1.834385 | -2.181848 |
| H | -1.194479 | -1.891134 | -0.444933 |
| C | 1.531476 | -2.099746 | -0.873362 |
| H | 0.989931 | -0.222165 | -1.798102 |
| H | 0.687134 | -0.281618 | -0.067437 |
| H | 1.160505 | -2.746858 | -0.066589 |
| H | 1.463475 | -2.690789 | -1.797285 |
| C | 2.991859 | -1.741568 | -0.606742 |
| C | 3.895782 | -2.966716 | -0.486589 |
| H | 3.059447 | -1.149169 | 0.316033 |
| H | 3.362321 | -1.095162 | -1.414215 |
| C | 5.354031 | -2.600638 | -0.222475 |
| H | 3.526153 | -3.612518 | 0.322173 |
| H | 3.826888 | -3.560717 | -1.408696 |
| H | 5.732351 | -1.959453 | -1.027054 |
| H | 5.432315 | -2.008276 | 0.696537 |
| C | 6.246204 | -3.828939 | -0.101895 |

| | | | |
|---|---|---|---|
| H | 6.212779 | -4.440171 | -1.012295 |
| H | 5.914672 | -4.487600 | 0.710333 |
| C | 7.687893 | -3.470007 | 0.157174 |
| O | 8.129531 | -2.353563 | 0.260900 |
| O | 8.464283 | -4.567722 | 0.265149 |
| H | 9.368805 | -4.247993 | 0.429500 |
| O | -3.803383 | 0.196593 | -0.127344 |
| C | -4.050567 | -1.131989 | -0.068640 |
| C | -3.925188 | -1.803239 | 1.202839 |
| C | -4.461802 | -1.787217 | -1.315422 |
| C | -3.754544 | -1.008904 | 2.359489 |
| C | -3.952505 | -3.208199 | 1.352073 |
| O | -4.730456 | -2.968494 | -1.437327 |
| O | -4.516541 | -0.922600 | -2.348079 |
| C | -3.611215 | -1.592159 | 3.608007 |
| H | -3.765291 | 0.071232 | 2.256047 |
| C | -3.805481 | -3.777707 | 2.608859 |
| H | -4.088613 | -3.831065 | 0.478395 |
| C | -4.907711 | -1.512996 | -3.581596 |
| C | -3.632572 | -2.981012 | 3.740636 |
| H | -3.489482 | -0.963186 | 4.484338 |
| H | -3.825170 | -4.858695 | 2.706697 |
| H | -5.899444 | -1.963345 | -3.496118 |
| H | -4.197405 | -2.288466 | -3.879660 |
| H | -4.916750 | -0.701312 | -4.307732 |
| H | -3.519912 | -3.437946 | 4.718655 |
| H | -1.921032 | 0.158099 | 0.743319 |

**5**

M06-2X SCF energy: -856.43302143 a.u.

M06-2X enthalpy: -855.905871 a.u.

M06-2X free energy: -855.984259 a.u.

M06-2X SCF energy in solution: -856.71863208 a.u.

M06-2X enthalpy in solution: -856.191482 a.u.

M06-2X free energy in solution: -856.26987 a.u.

Cartesian coordinates

| ATOM | X | Y | Z |
|---|---|---|---|
| C | 0.510451 | -0.109990 | 0.455765 |
| C | 1.489775 | 0.002077 | -0.439726 |
| H | 0.608857 | 0.410870 | 1.410701 |

| | | | |
|---|---|---|---|
| C | 2.761738 | 0.772693 | -0.235406 |
| C | 4.004430 | -0.123817 | -0.284572 |
| H | 2.724491 | 1.296313 | 0.728175 |
| H | 2.856493 | 1.545806 | -1.011193 |
| C | 5.306055 | 0.656538 | -0.116376 |
| H | 3.925518 | -0.887369 | 0.500028 |
| H | 4.024393 | -0.665215 | -1.240515 |
| H | 5.377218 | 1.419635 | -0.904148 |
| H | 5.282079 | 1.202238 | 0.837281 |
| C | 6.547633 | -0.231722 | -0.158829 |
| C | 7.850002 | 0.547758 | 0.010735 |
| H | 6.571966 | -0.777227 | -1.112546 |
| H | 6.475415 | -0.994819 | 0.628599 |
| C | 9.091876 | -0.339977 | -0.028949 |
| H | 7.922712 | 1.310051 | -0.777611 |
| H | 7.824591 | 1.094549 | 0.963740 |
| H | 9.020033 | -1.102044 | 0.759840 |
| H | 9.117896 | -0.887687 | -0.981549 |
| C | 10.394726 | 0.439241 | 0.140087 |
| C | 11.626961 | -0.460997 | 0.100804 |
| H | 10.366262 | 0.986647 | 1.091041 |
| H | 10.465427 | 1.198540 | -0.649372 |
| H | 11.588938 | -1.208816 | 0.900014 |
| H | 11.687458 | -0.997689 | -0.851952 |
| H | 12.550062 | 0.113083 | 0.221861 |
| C | -0.761517 | -0.880632 | 0.251086 |
| C | -2.003909 | 0.016159 | 0.301231 |
| H | -0.856351 | -1.654463 | 1.026191 |
| H | -0.724436 | -1.403307 | -0.713041 |
| C | -3.305584 | -0.763711 | 0.131209 |
| H | -2.024225 | 0.555913 | 1.258021 |
| H | -1.924654 | 0.780930 | -0.482056 |
| H | -3.282102 | -1.306314 | -0.824236 |
| H | -3.376579 | -1.529104 | 0.916776 |
| C | -4.546165 | 0.125590 | 0.177200 |
| C | -5.848264 | -0.653294 | 0.003770 |
| H | -4.474405 | 0.891802 | -0.606868 |
| H | -4.571541 | 0.666685 | 1.133092 |
| C | -7.083637 | 0.242207 | 0.050843 |
| H | -5.822866 | -1.194503 | -0.952491 |
| H | -5.920096 | -1.420229 | 0.787776 |

| | | | |
|---|---|---|---|
| H | -7.119324 | 0.783714 | 1.003248 |
| H | -7.021500 | 1.009509 | -0.729519 |
| C | -8.375441 | -0.544973 | -0.124566 |
| H | -8.486727 | -1.311839 | 0.652045 |
| H | -8.389634 | -1.085179 | -1.079317 |
| C | -9.598434 | 0.336142 | -0.077938 |
| O | -9.602315 | 1.531041 | 0.078100 |
| O | -10.733321 | -0.376151 | -0.234742 |
| H | -11.465675 | 0.263558 | -0.192462 |
| H | 1.391212 | -0.518912 | -1.394622 |

**6**

M06-2X SCF energy: -1429.65411120 a.u.

M06-2X enthalpy: -1428.955538 a.u.

M06-2X free energy: -1429.054349 a.u.

M06-2X SCF energy in solution: -1430.11822069 a.u.

M06-2X enthalpy in solution: -1429.419647 a.u.

M06-2X free energy in solution: -1429.518459 a.u.

Cartesian coordinates

| ATOM | X | Y | Z |
|---|---|---|---|
| C | -0.521321 | -1.380004 | -1.265974 |
| C | -2.045220 | -1.202638 | -1.040738 |
| H | -0.313513 | -1.775313 | -2.271400 |
| H | -2.647547 | -1.223435 | -1.954138 |
| C | -2.500680 | -0.069389 | -0.134539 |
| C | -2.513624 | 1.291903 | -0.829632 |
| H | -1.871220 | -0.034745 | 0.764406 |
| H | -3.513294 | -0.296120 | 0.223492 |
| C | -2.997496 | 2.413026 | 0.087976 |
| H | -1.510680 | 1.538256 | -1.200179 |
| H | -3.163515 | 1.239185 | -1.714389 |
| H | -4.002473 | 2.170027 | 0.460016 |
| H | -2.347327 | 2.462812 | 0.972446 |
| C | -3.026182 | 3.777755 | -0.596671 |
| C | -3.511126 | 4.899007 | 0.319884 |
| H | -3.674199 | 3.726950 | -1.482837 |
| H | -2.020063 | 4.020301 | -0.966447 |
| C | -3.535654 | 6.264955 | -0.362358 |
| H | -4.518405 | 4.657858 | 0.687265 |
| H | -2.865065 | 4.947532 | 1.207560 |

| | | | |
|---|---|---|---|
| H | -2.527938 | 6.507445 | -0.728171 |
| H | -4.180182 | 6.216785 | -1.251390 |
| C | -4.022969 | 7.386444 | 0.553062 |
| C | -4.039858 | 8.745762 | -0.141423 |
| H | -3.379279 | 7.432071 | 1.440927 |
| H | -5.029951 | 7.143341 | 0.915848 |
| H | -3.037211 | 9.019762 | -0.486426 |
| H | -4.697751 | 8.728168 | -1.016817 |
| H | -4.392317 | 9.536105 | 0.527436 |
| C | 0.484015 | -0.301603 | -0.924761 |
| C | 1.902478 | -0.869981 | -0.860578 |
| H | 0.226986 | 0.156516 | 0.038442 |
| H | 0.435246 | 0.486846 | -1.687058 |
| C | 2.943909 | 0.187049 | -0.500660 |
| H | 1.926058 | -1.682755 | -0.125094 |
| H | 2.157400 | -1.321047 | -1.829339 |
| H | 2.904388 | 1.005120 | -1.233828 |
| H | 2.688877 | 0.633124 | 0.470819 |
| C | 4.364132 | -0.371230 | -0.443086 |
| C | 5.406482 | 0.683084 | -0.077037 |
| H | 4.621055 | -0.814030 | -1.415101 |
| H | 4.403623 | -1.191008 | 0.286980 |
| C | 6.824997 | 0.121140 | -0.028559 |
| H | 5.362233 | 1.505679 | -0.804738 |
| H | 5.151804 | 1.122627 | 0.897634 |
| H | 6.879643 | -0.699811 | 0.695706 |
| H | 7.089755 | -0.315167 | -0.998610 |
| C | 7.853970 | 1.181726 | 0.339680 |
| H | 7.638820 | 1.626921 | 1.318988 |
| H | 7.844024 | 2.015088 | -0.373833 |
| C | 9.257983 | 0.633058 | 0.383626 |
| O | 9.578545 | -0.506170 | 0.157022 |
| O | 10.155464 | 1.584322 | 0.715178 |
| H | 11.026006 | 1.149010 | 0.719727 |
| O | -0.478758 | -2.467420 | -0.317341 |
| C | -1.895807 | -2.591039 | -0.341103 |
| C | -2.486070 | -2.794111 | 1.038475 |
| C | -2.278772 | -3.730456 | -1.290928 |
| C | -3.868845 | -2.829281 | 1.244990 |
| C | -1.626220 | -2.921739 | 2.129230 |
| O | -1.514586 | -4.535566 | -1.749878 |

| | | | |
|---|---|---|---|
| O | -3.592443 | -3.697849 | -1.583669 |
| C | -4.379394 | -2.992655 | 2.528690 |
| H | -4.542072 | -2.729941 | 0.399808 |
| C | -2.142154 | -3.094499 | 3.411616 |
| H | -0.556141 | -2.884223 | 1.960041 |
| C | -4.036578 | -4.744794 | -2.446526 |
| C | -3.518178 | -3.129244 | 3.615520 |
| H | -5.454251 | -3.013924 | 2.679943 |
| H | -1.464322 | -3.199748 | 4.253095 |
| H | -3.841402 | -5.718398 | -1.992292 |
| H | -3.517252 | -4.691152 | -3.405661 |
| H | -5.105995 | -4.588648 | -2.578107 |
| H | -3.919239 | -3.260086 | 4.615705 |

**6a**

M06-2X SCF energy: -1429.65450522 a.u.

M06-2X enthalpy: -1428.955927 a.u.

M06-2X free energy: -1429.054029 a.u.

M06-2X SCF energy in solution: -1430.11816431 a.u.

M06-2X enthalpy in solution: -1429.419586 a.u.

M06-2X free energy in solution: -1429.517688 a.u.

Cartesian coordinates

| ATOM | X | Y | Z |
|---|---|---|---|
| C | 0.774055 | 1.562997 | -0.285845 |
| C | 2.141777 | 1.923620 | 0.341558 |
| H | 0.716107 | 1.679378 | -1.372312 |
| H | 2.761475 | 2.496122 | -0.365257 |
| C | 2.982121 | 0.869994 | 1.026538 |
| C | 3.750438 | 0.007414 | 0.025439 |
| H | 2.346531 | 0.248225 | 1.667158 |
| H | 3.686295 | 1.386487 | 1.691321 |
| C | 4.568789 | -1.093144 | 0.697045 |
| H | 3.050648 | -0.447979 | -0.689805 |
| H | 4.417942 | 0.646717 | -0.568537 |
| H | 5.266048 | -0.638778 | 1.414414 |
| H | 3.899079 | -1.735571 | 1.285222 |
| C | 5.349565 | -1.949703 | -0.297233 |
| C | 6.173000 | -3.047570 | 0.372857 |
| H | 6.015527 | -1.305366 | -0.887977 |
| H | 4.650343 | -2.405131 | -1.012536 |

| | | | |
|---|---|---|---|
| C | 6.950217 | -3.907016 | -0.621762 |
| H | 6.874447 | -2.591636 | 1.085439 |
| H | 5.507851 | -3.690054 | 0.966595 |
| H | 6.248683 | -4.365455 | -1.333002 |
| H | 7.613498 | -3.264508 | -1.217843 |
| C | 7.777764 | -5.002828 | 0.047190 |
| C | 8.547349 | -5.854426 | -0.959247 |
| H | 7.114146 | -5.642695 | 0.642940 |
| H | 8.478586 | -4.543022 | 0.755805 |
| H | 7.862597 | -6.344379 | -1.659629 |
| H | 9.235701 | -5.237330 | -1.546408 |
| H | 9.134131 | -6.632960 | -0.463446 |
| C | 0.153098 | 0.243454 | 0.153195 |
| C | -1.244980 | 0.031832 | -0.431398 |
| H | 0.104908 | 0.206518 | 1.247393 |
| H | 0.803996 | -0.582993 | -0.161863 |
| C | -1.956749 | -1.180271 | 0.164523 |
| H | -1.861070 | 0.925774 | -0.263838 |
| H | -1.169575 | -0.080791 | -1.522060 |
| H | -1.344783 | -2.079364 | 0.006856 |
| H | -2.034781 | -1.047305 | 1.252752 |
| C | -3.348776 | -1.402159 | -0.422254 |
| C | -4.069087 | -2.606699 | 0.179621 |
| H | -3.270791 | -1.532138 | -1.510439 |
| H | -3.957280 | -0.500096 | -0.265215 |
| C | -5.460401 | -2.818109 | -0.411689 |
| H | -3.461363 | -3.508703 | 0.022063 |
| H | -4.147283 | -2.476584 | 1.268070 |
| H | -6.074647 | -1.923441 | -0.256937 |
| H | -5.391039 | -2.951130 | -1.497551 |
| C | -6.170572 | -4.021413 | 0.194046 |
| H | -6.280792 | -3.917414 | 1.280597 |
| H | -5.598239 | -4.944643 | 0.040652 |
| C | -7.545668 | -4.230974 | -0.388788 |
| O | -8.065038 | -3.547424 | -1.234554 |
| O | -8.159962 | -5.302342 | 0.153291 |
| H | -9.031925 | -5.363804 | -0.274846 |
| O | 1.511363 | 2.826693 | 1.270747 |
| C | 0.312813 | 2.840419 | 0.497290 |
| C | 0.208575 | 4.081744 | -0.369120 |
| C | -0.889313 | 2.636120 | 1.422159 |

| | | | |
|---|---|---|---|
| C | -0.640760 | 4.111801 | -1.477817 |
| C | 0.986783 | 5.197882 | -0.070276 |
| O | -0.827498 | 2.092397 | 2.494219 |
| O | -2.036738 | 3.071383 | 0.878665 |
| C | -0.716250 | 5.249657 | -2.272403 |
| H | -1.247884 | 3.240970 | -1.712724 |
| C | 0.909794 | 6.338818 | -0.867234 |
| H | 1.655190 | 5.155773 | 0.783209 |
| C | -3.203891 | 2.781144 | 1.645363 |
| C | 0.059551 | 6.367777 | -1.967962 |
| H | -1.378300 | 5.264271 | -3.132686 |
| H | 1.519708 | 7.204519 | -0.627916 |
| H | -3.132615 | 3.238069 | 2.634120 |
| H | -3.317756 | 1.699357 | 1.762571 |
| H | -4.038530 | 3.198129 | 1.084639 |
| H | 0.002745 | 7.255100 | -2.590716 |

**TS-1**

M06-2X SCF energy: -1429.54027542 a.u.

M06-2X enthalpy: -1428.846934 a.u.

M06-2X free energy: -1428.947574 a.u.

M06-2X SCF energy in solution: -1430.01024874 a.u.

M06-2X enthalpy in solution: -1429.316907 a.u.

M06-2X free energy in solution: -1429.417548 a.u.

Imaginary frequency: -85.3845 cm-1

Cartesian coordinates

| ATOM | X | Y | Z |
|---|---|---|---|
| C | -0.062836 | 0.477094 | 2.858408 |
| C | 1.298841 | 0.347031 | 2.834430 |
| H | -0.464771 | 1.360280 | 3.346727 |
| C | 2.089639 | -0.849284 | 2.421245 |
| C | 3.304600 | -0.507863 | 1.550173 |
| H | 1.455537 | -1.583794 | 1.915728 |
| H | 2.454421 | -1.336375 | 3.340689 |
| C | 4.148056 | -1.738129 | 1.222328 |
| H | 2.978072 | -0.037184 | 0.616399 |
| H | 3.926180 | 0.235296 | 2.067282 |
| H | 4.570012 | -2.154984 | 2.147580 |
| H | 3.502003 | -2.522683 | 0.801868 |
| C | 5.270359 | -1.430646 | 0.233590 |

| | | | |
|---|---|---|---|
| C | 6.129659 | -2.648998 | -0.096519 |
| H | 5.907986 | -0.634957 | 0.643092 |
| H | 4.832160 | -1.027434 | -0.690014 |
| C | 7.247517 | -2.342515 | -1.090740 |
| H | 6.566187 | -3.049149 | 0.829457 |
| H | 5.490646 | -3.445557 | -0.503105 |
| H | 6.811152 | -1.944032 | -2.017161 |
| H | 7.886014 | -1.544819 | -0.685883 |
| C | 8.109350 | -3.559631 | -1.420759 |
| C | 9.222056 | -3.238531 | -2.415361 |
| H | 7.470005 | -4.355213 | -1.824671 |
| H | 8.544068 | -3.955607 | -0.493865 |
| H | 8.806762 | -2.867486 | -3.358295 |
| H | 9.887136 | -2.463842 | -2.019272 |
| H | 9.829455 | -4.120058 | -2.639656 |
| C | -1.072669 | -0.568160 | 2.507308 |
| C | -2.270832 | -0.004236 | 1.738482 |
| H | -1.433999 | -1.003111 | 3.452772 |
| H | -0.615830 | -1.382424 | 1.939059 |
| C | -3.319910 | -1.071838 | 1.440183 |
| H | -2.724589 | 0.811444 | 2.317612 |
| H | -1.906454 | 0.438094 | 0.806395 |
| H | -2.858834 | -1.876645 | 0.849480 |
| H | -3.658344 | -1.534225 | 2.378443 |
| C | -4.526704 | -0.520687 | 0.683981 |
| C | -5.569635 | -1.588501 | 0.361228 |
| H | -4.187048 | -0.049516 | -0.248147 |
| H | -4.994566 | 0.276555 | 1.277579 |
| C | -6.778011 | -1.028203 | -0.384563 |
| H | -5.102960 | -2.382174 | -0.239114 |
| H | -5.902662 | -2.065640 | 1.293785 |
| H | -7.254566 | -0.240412 | 0.210187 |
| H | -6.455579 | -0.548698 | -1.316011 |
| C | -7.807682 | -2.103242 | -0.706086 |
| H | -8.170024 | -2.596650 | 0.204401 |
| H | -7.376467 | -2.900349 | -1.324257 |
| C | -9.006762 | -1.553217 | -1.436896 |
| O | -9.178492 | -0.401033 | -1.744861 |
| O | -9.904136 | -2.519469 | -1.721795 |
| H | -10.638194 | -2.081829 | -2.187629 |
| O | 0.020933 | 2.078896 | 1.288579 |

| | | | |
|---|---|---|---|
| C | 0.688886 | 2.065134 | 0.157482 |
| C | 0.987711 | 3.385456 | -0.398497 |
| C | 1.013031 | 0.837942 | -0.549465 |
| C | 0.854436 | 4.508082 | 0.443424 |
| C | 1.370349 | 3.592270 | -1.738877 |
| O | 1.753945 | 0.741356 | -1.516292 |
| O | 0.392158 | -0.254290 | -0.023346 |
| C | 1.104627 | 5.786744 | -0.032785 |
| H | 0.561388 | 4.356350 | 1.475978 |
| C | 1.625034 | 4.876697 | -2.199761 |
| H | 1.474400 | 2.744937 | -2.402734 |
| C | 0.684039 | -1.466441 | -0.703632 |
| C | 1.494083 | 5.980080 | -1.357011 |
| H | 1.001048 | 6.637021 | 0.634443 |
| H | 1.924725 | 5.017889 | -3.233756 |
| H | 1.753870 | -1.694232 | -0.652291 |
| H | 0.393960 | -1.401808 | -1.754841 |
| H | 0.105390 | -2.240342 | -0.195989 |
| H | 1.694040 | 6.979968 | -1.729026 |
| H | 1.879632 | 1.201263 | 3.180621 |

**TS-2**

M06-2X SCF energy: -1429.54017180 a.u.

M06-2X enthalpy: -1428.846796 a.u.

M06-2X free energy: -1428.947435 a.u.

M06-2X SCF energy in solution: -1430.01016348 a.u.

M06-2X enthalpy in solution: -1429.316788 a.u.

M06-2X free energy in solution: -1429.417426 a.u.

Imaginary frequency: -84.0797 cm-1

Cartesian coordinates

| ATOM | X | Y | Z |
|---|---|---|---|
| C | 0.222905 | 0.483600 | 2.893121 |
| C | 1.582017 | 0.406911 | 2.758681 |
| H | -0.179856 | 1.416444 | 3.286181 |
| C | 2.377397 | -0.778501 | 2.314623 |
| C | 3.545707 | -0.403941 | 1.398043 |
| H | 1.740984 | -1.518331 | 1.822326 |
| H | 2.780919 | -1.258225 | 3.220542 |
| C | 4.383426 | -1.617975 | 1.006324 |
| H | 3.142169 | 0.081786 | 0.504309 |

| | | | |
|---|---|---|---|
| H | 4.179823 | 0.338904 | 1.900811 |
| H | 4.760141 | -2.117583 | 1.910301 |
| H | 3.741739 | -2.350437 | 0.495430 |
| C | 5.559062 | -1.260934 | 0.099284 |
| C | 6.390217 | -2.474527 | -0.311315 |
| H | 6.204593 | -0.534072 | 0.611499 |
| H | 5.182249 | -0.755212 | -0.800170 |
| C | 7.570454 | -2.117916 | -1.212346 |
| H | 6.761565 | -2.983060 | 0.589560 |
| H | 5.744934 | -3.198997 | -0.827794 |
| H | 7.200308 | -1.607036 | -2.112255 |
| H | 8.217870 | -1.395673 | -0.695442 |
| C | 8.399614 | -3.331978 | -1.626648 |
| C | 9.576717 | -2.960515 | -2.524908 |
| H | 7.751623 | -4.051169 | -2.144033 |
| H | 8.767036 | -3.841616 | -0.726574 |
| H | 9.228728 | -2.473617 | -3.442063 |
| H | 10.250885 | -2.263636 | -2.015740 |
| H | 10.158507 | -3.841034 | -2.812177 |
| C | -0.780211 | -0.581394 | 2.597215 |
| C | -2.022662 | -0.062974 | 1.863186 |
| H | -1.104917 | -1.005657 | 3.561677 |
| H | -0.325434 | -1.404168 | 2.037499 |
| C | -3.068977 | -1.154988 | 1.650460 |
| H | -2.465836 | 0.762986 | 2.435335 |
| H | -1.738670 | 0.355457 | 0.891512 |
| H | -2.596972 | -2.023863 | 1.168833 |
| H | -3.438128 | -1.509270 | 2.623169 |
| C | -4.240101 | -0.681634 | 0.792628 |
| C | -5.292745 | -1.764852 | 0.568366 |
| H | -3.856824 | -0.334964 | -0.176969 |
| H | -4.710375 | 0.190759 | 1.266595 |
| C | -6.463140 | -1.283191 | -0.285156 |
| H | -4.821980 | -2.634775 | 0.088863 |
| H | -5.667357 | -2.116253 | 1.540228 |
| H | -6.945663 | -0.421546 | 0.190270 |
| H | -6.098541 | -0.926667 | -1.255377 |
| C | -7.500373 | -2.375380 | -0.510234 |
| H | -7.899078 | -2.751279 | 0.440415 |
| H | -7.064081 | -3.246385 | -1.014492 |
| C | -8.667487 | -1.900624 | -1.339002 |

| | | | |
|---|---|---|---|
| O | -8.818181 | -0.787089 | -1.773708 |
| O | -9.562800 | -2.887391 | -1.551372 |
| H | -10.276294 | -2.498444 | -2.087125 |
| O | 1.551986 | 2.000708 | 1.174935 |
| C | 0.761216 | 2.084493 | 0.129271 |
| C | 0.597276 | 3.433507 | -0.413673 |
| C | 0.179777 | 0.916982 | -0.510948 |
| C | 0.999483 | 4.524624 | 0.383117 |
| C | 0.087376 | 3.693577 | -1.701390 |
| O | -0.675292 | 0.928260 | -1.383753 |
| O | 0.690484 | -0.253515 | -0.037473 |
| C | 0.886734 | 5.825573 | -0.084800 |
| H | 1.391090 | 4.332224 | 1.375437 |
| C | -0.027384 | 5.001090 | -2.153492 |
| H | -0.223840 | 2.870744 | -2.330304 |
| C | 0.148836 | -1.411080 | -0.656755 |
| C | 0.370245 | 6.073413 | -1.355434 |
| H | 1.197611 | 6.651543 | 0.547685 |
| H | -0.428544 | 5.184288 | -3.145666 |
| H | 0.323821 | -1.393179 | -1.734920 |
| H | -0.930052 | -1.477758 | -0.481624 |
| H | 0.661748 | -2.260540 | -0.202094 |
| H | 0.277216 | 7.091455 | -1.720216 |
| H | 2.165036 | 1.221867 | 3.178216 |

**TS-3**

M06-2X SCF energy: -1429.54043241 a.u.

M06-2X enthalpy: -1428.847189 a.u.

M06-2X free energy: -1428.948982 a.u.

M06-2X SCF energy in solution: -1430.01010527 a.u.

M06-2X enthalpy in solution: -1429.316862 a.u.

M06-2X free energy in solution: -1429.418655 a.u.

Imaginary frequency: -113.5348 cm-1

Cartesian coordinates

| ATOM | X | Y | Z |
|---|---|---|---|
| C | -0.184645 | -1.275907 | -0.414799 |
| C | -1.279926 | -1.030642 | 0.362406 |
| H | -0.300173 | -1.197855 | -1.494271 |
| C | -2.588738 | -0.545111 | -0.161591 |
| C | -3.745572 | -1.494798 | 0.182758 |

| | | | |
|---|---|---|---|
| H | -2.523336 | -0.402030 | -1.246440 |
| H | -2.797932 | 0.442414 | 0.276964 |
| C | -5.091421 | -0.975581 | -0.318149 |
| H | -3.546568 | -2.483569 | -0.250275 |
| H | -3.789089 | -1.634423 | 1.271174 |
| H | -5.280161 | 0.016872 | 0.113154 |
| H | -5.044081 | -0.834153 | -1.406639 |
| C | -6.252122 | -1.907839 | 0.022116 |
| C | -7.599191 | -1.394604 | -0.482470 |
| H | -6.300382 | -2.045717 | 1.111218 |
| H | -6.058535 | -2.901741 | -0.404971 |
| C | -8.761681 | -2.323723 | -0.140049 |
| H | -7.791276 | -0.399647 | -0.057471 |
| H | -7.551197 | -1.258413 | -1.571877 |
| H | -8.569016 | -3.319701 | -0.563239 |
| H | -8.811125 | -2.458756 | 0.949591 |
| C | -10.109211 | -1.812899 | -0.646354 |
| C | -11.261537 | -2.751031 | -0.296537 |
| H | -10.057999 | -1.678922 | -1.734532 |
| H | -10.299672 | -0.818350 | -0.222984 |
| H | -11.103163 | -3.742860 | -0.733172 |
| H | -11.346589 | -2.876523 | 0.788081 |
| H | -12.217821 | -2.370242 | -0.666321 |
| C | 1.100979 | -1.862366 | 0.070946 |
| C | 2.339481 | -1.127196 | -0.449922 |
| H | 1.138899 | -2.905185 | -0.281287 |
| H | 1.108752 | -1.893174 | 1.165131 |
| C | 3.636799 | -1.786933 | 0.009592 |
| H | 2.308113 | -1.095338 | -1.547517 |
| H | 2.295880 | -0.090385 | -0.103757 |
| H | 3.662232 | -1.809182 | 1.108749 |
| H | 3.658524 | -2.835818 | -0.319186 |
| C | 4.881547 | -1.069274 | -0.507709 |
| C | 6.183119 | -1.713042 | -0.034915 |
| H | 4.855816 | -0.019633 | -0.185608 |
| H | 4.861979 | -1.053451 | -1.606004 |
| C | 7.423323 | -0.997304 | -0.564044 |
| H | 6.205243 | -1.722557 | 1.063971 |
| H | 6.204091 | -2.765707 | -0.350638 |
| H | 7.413639 | -0.989137 | -1.660082 |
| H | 7.410898 | 0.054315 | -0.255160 |

| | | | |
|---|---|---|---|
| C | 8.714145 | -1.644993 | -0.080797 |
| H | 8.773984 | -2.697056 | -0.386268 |
| H | 8.775266 | -1.649923 | 1.014523 |
| C | 9.942432 | -0.944962 | -0.605848 |
| O | 9.950912 | 0.015854 | -1.332931 |
| O | 11.076191 | -1.526303 | -0.162301 |
| H | 11.812248 | -1.016838 | -0.544471 |
| O | 0.016486 | 0.935247 | -0.674581 |
| C | -0.290575 | 1.788683 | 0.280267 |
| C | -0.891363 | 3.038826 | -0.170263 |
| C | 0.019282 | 1.539219 | 1.680936 |
| C | -1.412905 | 3.091327 | -1.480311 |
| C | -0.954786 | 4.201476 | 0.626061 |
| O | -0.374477 | 2.197868 | 2.629909 |
| O | 0.846190 | 0.470856 | 1.850079 |
| C | -1.985964 | 4.255815 | -1.969960 |
| H | -1.365880 | 2.200667 | -2.096681 |
| C | -1.538062 | 5.356375 | 0.125732 |
| H | -0.555818 | 4.182658 | 1.631330 |
| C | 1.184941 | 0.204887 | 3.203823 |
| C | -2.054952 | 5.395244 | -1.170102 |
| H | -2.386657 | 4.273631 | -2.978941 |
| H | -1.584670 | 6.241193 | 0.753243 |
| H | 0.293296 | -0.038975 | 3.788429 |
| H | 1.671423 | 1.070396 | 3.659335 |
| H | 1.867769 | -0.645447 | 3.177242 |
| H | -2.506866 | 6.305452 | -1.551434 |
| H | -1.192849 | -1.166932 | 1.440807 |

**TS-4**

M06-2X SCF energy: -1429.54052555 a.u.

M06-2X enthalpy: -1428.847291 a.u.

M06-2X free energy: -1428.94914 a.u.

M06-2X SCF energy in solution: -1430.01017712 a.u.

M06-2X enthalpy in solution: -1429.316943 a.u.

M06-2X free energy in solution: -1429.418791 a.u.

Imaginary frequency: -111.8938 cm-1

Cartesian coordinates

| ATOM | X | Y | Z |
|---|---|---|---|
| C | 0.055299 | -0.893428 | 0.345670 |

| | | | |
|---|---|---|---|
| C | -0.992599 | -1.231138 | -0.461507 |
| H | -0.047297 | -1.045558 | 1.420581 |
| C | -2.232649 | -1.931949 | -0.010606 |
| C | -3.518179 | -1.309631 | -0.563510 |
| H | -2.266294 | -1.966024 | 1.083026 |
| H | -2.167640 | -2.973186 | -0.363526 |
| C | -4.763525 | -2.077545 | -0.128967 |
| H | -3.572533 | -0.271514 | -0.222842 |
| H | -3.464541 | -1.279778 | -1.660360 |
| H | -4.686992 | -3.125676 | -0.451911 |
| H | -4.811312 | -2.097859 | 0.969488 |
| C | -6.054554 | -1.472917 | -0.676554 |
| C | -7.306628 | -2.223622 | -0.228110 |
| H | -6.011634 | -1.459798 | -1.774483 |
| H | -6.125309 | -0.423251 | -0.360401 |
| C | -8.597390 | -1.623448 | -0.780918 |
| H | -7.232127 | -3.275031 | -0.539700 |
| H | -7.351337 | -2.232253 | 0.870040 |
| H | -8.671024 | -0.571191 | -0.472296 |
| H | -8.554914 | -1.617452 | -1.879172 |
| C | -9.850719 | -2.370475 | -0.328911 |
| C | -11.132443 | -1.760434 | -0.890514 |
| H | -9.892046 | -2.374039 | 0.767948 |
| H | -9.774769 | -3.421328 | -0.636870 |
| H | -11.240112 | -0.718191 | -0.571905 |
| H | -11.122455 | -1.772249 | -1.985577 |
| H | -12.019282 | -2.306843 | -0.556942 |
| C | 1.329007 | -0.289145 | -0.140073 |
| C | 2.552398 | -1.153608 | 0.199251 |
| H | 1.450577 | 0.698466 | 0.329805 |
| H | 1.270147 | -0.120529 | -1.221602 |
| C | 3.861873 | -0.504275 | -0.242989 |
| H | 2.581408 | -1.334211 | 1.282049 |
| H | 2.447023 | -2.136481 | -0.277689 |
| H | 3.825576 | -0.311482 | -1.323984 |
| H | 3.961053 | 0.476448 | 0.241693 |
| C | 5.085709 | -1.358685 | 0.079434 |
| C | 6.398220 | -0.707632 | -0.351702 |
| H | 4.987489 | -2.337019 | -0.410756 |
| H | 5.117423 | -1.557751 | 1.159405 |
| C | 7.616401 | -1.570471 | -0.032834 |

| | | | |
|---|---|---|---|
| H | 6.364962 | -0.502628 | -1.430904 |
| H | 6.498610 | 0.268505 | 0.142884 |
| H | 7.656605 | -1.781565 | 1.042059 |
| H | 7.527247 | -2.544429 | -0.527848 |
| C | 8.920567 | -0.910785 | -0.460734 |
| H | 9.061397 | 0.058258 | 0.033759 |
| H | 8.929247 | -0.699167 | -1.537167 |
| C | 10.124819 | -1.764235 | -0.150957 |
| O | 10.104247 | -2.849824 | 0.371837 |
| O | 11.271887 | -1.166834 | -0.533303 |
| H | 11.991079 | -1.779431 | -0.299039 |
| O | -1.379441 | 0.957609 | -0.723822 |
| C | -1.167644 | 1.829436 | 0.240165 |
| C | -0.663270 | 3.127732 | -0.191466 |
| C | -1.488741 | 1.546735 | 1.632377 |
| C | -0.124993 | 3.233129 | -1.491563 |
| C | -0.708073 | 4.285338 | 0.613605 |
| O | -1.170373 | 2.228449 | 2.593220 |
| O | -2.231015 | 0.415056 | 1.776444 |
| C | 0.360336 | 4.444106 | -1.962963 |
| H | -0.089458 | 2.346700 | -2.114792 |
| C | -0.211334 | 5.487605 | 0.131653 |
| H | -1.121357 | 4.226228 | 1.611487 |
| C | -2.577580 | 0.111887 | 3.120468 |
| C | 0.323263 | 5.578856 | -1.154241 |
| H | 0.775385 | 4.502445 | -2.964598 |
| H | -0.247072 | 6.368018 | 0.766049 |
| H | -3.138727 | 0.933531 | 3.571224 |
| H | -1.682526 | -0.068177 | 3.722806 |
| H | -3.192316 | -0.787902 | 3.072462 |
| H | 0.706855 | 6.525638 | -1.521153 |
| H | -0.856868 | -1.136308 | -1.537293 |

**FA 18:2 (9*Z*, 12*Z*)**

M06-2X SCF energy: -855.20721360 a.u.

M06-2X enthalpy: -854.704428 a.u.

M06-2X free energy: -854.779587 a.u.

M06-2X SCF energy in solution: -855.49405290 a.u.

M06-2X enthalpy in solution: -854.991267 a.u.

M06-2X free energy in solution: -855.066426 a.u.

Cartesian coordinates

| ATOM | X | Y | Z |
|---|---|---|---|
| C | -3.904514 | -2.178484 | -0.610073 |
| C | -5.006049 | -1.463087 | -0.369464 |
| H | -3.865274 | -2.739996 | -1.544306 |
| C | -5.299993 | -0.591868 | 0.827589 |
| C | -5.554293 | 0.848927 | 0.426688 |
| H | -4.479175 | -0.640984 | 1.550725 |
| H | -6.185829 | -0.981736 | 1.342172 |
| C | -4.620337 | 1.666622 | -0.061683 |
| H | -6.573576 | 1.216711 | 0.521916 |
| H | -4.906049 | 2.679464 | -0.346653 |
| C | -3.180500 | 1.296510 | -0.277103 |
| C | -2.194269 | 2.430127 | 0.007382 |
| H | -3.045753 | 0.959596 | -1.316643 |
| H | -2.920911 | 0.433434 | 0.347544 |
| C | -0.748937 | 1.984402 | -0.202562 |
| H | -2.415441 | 3.286587 | -0.644297 |
| H | -2.326152 | 2.782182 | 1.038960 |
| H | -0.538149 | 1.126180 | 0.454123 |
| H | -0.630961 | 1.613877 | -1.231945 |
| C | 0.283881 | 3.077632 | 0.058724 |
| C | 1.713073 | 2.579575 | -0.142405 |
| H | 0.162246 | 3.453659 | 1.082524 |
| H | 0.089450 | 3.927460 | -0.607812 |
| H | 1.931997 | 1.747185 | 0.537436 |
| H | 1.858813 | 2.216860 | -1.166277 |
| H | 2.449206 | 3.367164 | 0.042450 |
| C | -2.670750 | -2.291177 | 0.241052 |
| C | -1.402519 | -1.893433 | -0.526367 |
| H | -2.558396 | -3.326529 | 0.592812 |
| H | -2.757414 | -1.673180 | 1.141822 |
| C | -0.163788 | -1.842082 | 0.364061 |
| H | -1.238430 | -2.598556 | -1.352457 |
| H | -1.552756 | -0.909094 | -0.988502 |
| H | -0.351431 | -1.147536 | 1.196427 |
| H | 0.006800 | -2.826433 | 0.822180 |
| C | 1.091362 | -1.398813 | -0.384024 |
| C | 2.306650 | -1.242371 | 0.526462 |
| H | 0.894685 | -0.439151 | -0.883741 |
| H | 1.319065 | -2.119791 | -1.181061 |

| | | | |
|---|---|---|---|
| C | 3.558154 | -0.806038 | -0.230563 |
| H | 2.073914 | -0.506828 | 1.310592 |
| H | 2.502109 | -2.190056 | 1.047716 |
| H | 3.835057 | -1.566364 | -0.969807 |
| H | 3.351485 | 0.107529 | -0.800588 |
| C | 4.737708 | -0.554958 | 0.698867 |
| H | 4.977541 | -1.444502 | 1.294718 |
| H | 4.512794 | 0.237266 | 1.424055 |
| C | 5.984759 | -0.152925 | -0.047654 |
| O | 6.087259 | -0.041980 | -1.243204 |
| O | 7.015888 | 0.078604 | 0.790888 |
| H | 7.772274 | 0.328613 | 0.231673 |
| H | -5.794730 | -1.477389 | -1.120786 |

**TS-5**

M06-2X SCF energy: -1428.31345009 a.u.

M06-2X enthalpy: -1427.644598 a.u.

M06-2X free energy: -1427.743933 a.u.

M06-2X SCF energy in solution: -1428.78431657 a.u.

M06-2X enthalpy in solution: -1428.115464 a.u.

M06-2X free energy in solution: -1428.2148 a.u.

Imaginary frequency: -36.8433 cm-1

Cartesian coordinates

| ATOM | X | Y | Z |
|---|---|---|---|
| C | 1.436897 | 0.321954 | 0.966092 |
| C | 2.157238 | 1.368697 | 0.469624 |
| H | 1.553735 | 0.106637 | 2.025248 |
| C | 2.211656 | 1.841257 | -0.950503 |
| C | 1.859149 | 3.314086 | -1.056483 |
| H | 1.560022 | 1.238303 | -1.590069 |
| H | 3.235352 | 1.679766 | -1.311267 |
| C | 0.644676 | 3.806395 | -0.807006 |
| H | 2.661044 | 3.995899 | -1.327569 |
| H | 0.488832 | 4.882047 | -0.888977 |
| C | -0.550013 | 2.989462 | -0.403553 |
| C | -1.870149 | 3.478497 | -1.001622 |
| H | -0.637612 | 2.990044 | 0.694231 |
| H | -0.397588 | 1.941662 | -0.690643 |
| C | -3.038914 | 2.584747 | -0.593220 |
| H | -2.062193 | 4.510632 | -0.678533 |

| | | | |
|---|---|---|---|
| H | -1.789339 | 3.504334 | -2.096060 |
| H | -2.839365 | 1.555477 | -0.929190 |
| H | -3.094373 | 2.540159 | 0.504910 |
| C | -4.388279 | 3.036614 | -1.145791 |
| C | -5.521396 | 2.099322 | -0.735375 |
| H | -4.330893 | 3.093104 | -2.240186 |
| H | -4.601741 | 4.053900 | -0.794188 |
| H | -5.337068 | 1.083580 | -1.105740 |
| H | -5.605527 | 2.043481 | 0.355892 |
| H | -6.486332 | 2.430576 | -1.129435 |
| C | 0.421345 | -0.502626 | 0.242613 |
| C | -0.961045 | -0.414831 | 0.906928 |
| H | 0.764394 | -1.544861 | 0.249824 |
| H | 0.347298 | -0.208213 | -0.809409 |
| C | -2.039327 | -1.130694 | 0.097326 |
| H | -0.908648 | -0.841492 | 1.917202 |
| H | -1.247679 | 0.637833 | 1.030210 |
| H | -2.065163 | -0.706482 | -0.917255 |
| H | -1.771072 | -2.189414 | -0.021694 |
| C | -3.426830 | -1.017338 | 0.724349 |
| C | -4.521958 | -1.647334 | -0.132505 |
| H | -3.666314 | 0.043770 | 0.885932 |
| H | -3.421276 | -1.486575 | 1.717584 |
| C | -5.904808 | -1.533863 | 0.503478 |
| H | -4.528279 | -1.162232 | -1.119726 |
| H | -4.284918 | -2.704728 | -0.315387 |
| H | -5.923520 | -2.067470 | 1.460631 |
| H | -6.124376 | -0.485591 | 0.738340 |
| C | -7.002649 | -2.080788 | -0.398655 |
| H | -6.823783 | -3.131278 | -0.659666 |
| H | -7.042419 | -1.539468 | -1.352359 |
| C | -8.367395 | -1.990771 | 0.236888 |
| O | -8.607772 | -1.549138 | 1.332028 |
| O | -9.329311 | -2.473234 | -0.576038 |
| H | -10.169077 | -2.379190 | -0.092946 |
| H | 2.800966 | 1.902708 | 1.164815 |
| O | 3.196135 | -0.837899 | 0.100445 |
| C | 4.466372 | -0.700777 | 0.415798 |
| C | 5.403622 | -1.267503 | -0.553224 |
| C | 4.910847 | -0.101508 | 1.666862 |
| C | 4.920090 | -1.575994 | -1.841531 |

| | | | |
|---|---|---|---|
| C | 6.753340 | -1.546386 | -0.258354 |
| O | 6.066890 | 0.166727 | 1.951288 |
| O | 3.894537 | 0.126321 | 2.544488 |
| C | 5.755279 | -2.130754 | -2.800327 |
| H | 3.883176 | -1.363037 | -2.076486 |
| C | 7.579198 | -2.095011 | -1.229828 |
| H | 7.143314 | -1.321658 | 0.724841 |
| C | 4.314922 | 0.690096 | 3.780714 |
| C | 7.091446 | -2.392066 | -2.502302 |
| H | 5.363330 | -2.354691 | -3.787878 |
| H | 8.617870 | -2.298522 | -0.987530 |
| H | 4.803877 | 1.654704 | 3.622590 |
| H | 5.016497 | 0.025196 | 4.289732 |
| H | 3.408514 | 0.813627 | 4.373467 |
| H | 7.746619 | -2.822067 | -3.253293 |

**TS-6**

M06-2X SCF energy: -1428.31188565 a.u.

M06-2X enthalpy: -1427.648621 a.u.

M06-2X free energy: -1427.74611 a.u.

M06-2X SCF energy in solution: -1428.77776435 a.u.

M06-2X enthalpy in solution: -1428.114500 a.u.

M06-2X free energy in solution: -1428.211988 a.u.

Imaginary frequency: -1033.6498 cm-1

Cartesian coordinates

| ATOM | X | Y | Z |
|---|---|---|---|
| O | -2.889444 | 0.199132 | -1.627366 |
| C | -2.634802 | -0.943462 | -1.010833 |
| C | -1.344418 | -1.548741 | -1.282341 |
| C | -3.673483 | -1.585136 | -0.200642 |
| C | -0.444781 | -0.853236 | -2.120311 |
| C | -0.928398 | -2.778001 | -0.723393 |
| O | -3.499186 | -2.485531 | 0.601667 |
| O | -4.899210 | -1.079562 | -0.466209 |
| C | 0.830603 | -1.344291 | -2.357734 |
| H | -0.758153 | 0.084081 | -2.567400 |
| C | 0.355146 | -3.249586 | -0.961894 |
| H | -1.606833 | -3.335950 | -0.091484 |
| C | -5.944700 | -1.660348 | 0.303949 |
| C | 1.244363 | -2.540847 | -1.772676 |

| | | | |
|---|---|---|---|
| H | 1.509384 | -0.782753 | -2.993251 |
| H | 0.666584 | -4.185911 | -0.508235 |
| H | -5.730379 | -1.568830 | 1.372034 |
| H | -6.057777 | -2.719128 | 0.059464 |
| H | -6.847306 | -1.109460 | 0.040722 |
| H | 2.245809 | -2.920958 | -1.950051 |
| C | -2.440790 | 3.662730 | -0.327548 |
| C | -3.579511 | 3.151755 | 0.140963 |
| H | -2.259314 | 4.715628 | -0.106750 |
| C | -4.186347 | 1.774078 | 0.054801 |
| C | -4.269611 | 1.045337 | 1.344680 |
| H | -3.594434 | 1.074355 | -0.729572 |
| H | -5.173488 | 1.825460 | -0.417586 |
| C | -3.223887 | 0.649638 | 2.095082 |
| H | -5.269036 | 0.789227 | 1.692348 |
| H | -3.441020 | 0.142773 | 3.035851 |
| C | -1.772539 | 0.765511 | 1.744493 |
| C | -0.998724 | -0.503904 | 2.124975 |
| H | -1.339802 | 1.626256 | 2.277904 |
| H | -1.650011 | 0.972385 | 0.674959 |
| C | 0.451243 | -0.487250 | 1.647165 |
| H | -1.024146 | -0.620216 | 3.217834 |
| H | -1.511471 | -1.380084 | 1.707546 |
| H | 0.473394 | -0.508820 | 0.547454 |
| H | 0.934126 | 0.453882 | 1.952899 |
| C | 1.256733 | -1.668155 | 2.185511 |
| C | 2.679003 | -1.707522 | 1.634160 |
| H | 0.735940 | -2.599237 | 1.926545 |
| H | 1.282218 | -1.619154 | 3.281852 |
| H | 2.659461 | -1.852467 | 0.547921 |
| H | 3.203429 | -0.765783 | 1.838684 |
| H | 3.263424 | -2.521006 | 2.074803 |
| C | -1.339914 | 2.983987 | -1.086476 |
| C | 0.011569 | 3.125669 | -0.370360 |
| H | -1.258184 | 3.429506 | -2.088011 |
| H | -1.573439 | 1.926408 | -1.238730 |
| C | 1.077596 | 2.221957 | -0.980291 |
| H | 0.336841 | 4.174691 | -0.389802 |
| H | -0.111329 | 2.860563 | 0.687916 |
| H | 0.707025 | 1.189467 | -0.950574 |
| H | 1.215305 | 2.467442 | -2.043648 |

| | | | |
|---|---|---|---|
| C | 2.421304 | 2.270473 | -0.258537 |
| C | 3.366665 | 1.170851 | -0.736301 |
| H | 2.254189 | 2.147619 | 0.821044 |
| H | 2.890210 | 3.255396 | -0.388205 |
| C | 4.700103 | 1.160634 | 0.005053 |
| H | 2.866110 | 0.200003 | -0.605988 |
| H | 3.544767 | 1.280770 | -1.816003 |
| H | 5.245343 | 2.093814 | -0.177906 |
| H | 4.526785 | 1.121999 | 1.087366 |
| C | 5.570094 | -0.021649 | -0.399477 |
| H | 5.773140 | -0.021666 | -1.477748 |
| H | 5.065746 | -0.973950 | -0.190139 |
| C | 6.896065 | -0.036850 | 0.317872 |
| O | 7.270889 | 0.773347 | 1.127469 |
| O | 7.652252 | -1.091499 | -0.051747 |
| H | 8.480637 | -1.024099 | 0.454762 |
| H | -4.227649 | 3.828797 | 0.701322 |
